# High Throughput Screening Identifies a Small Molecule Trafficking Corrector for Long-QT Syndrome Associated KCNQ1 Variants

**DOI:** 10.1101/2025.10.13.682066

**Authors:** Katherine R. Clowes Moster, Carlos G. Vanoye, Ana C. Chang-Gonzalez, Ian M. Romaine, Katherine M. Stefanski, Mason C. Wilkinson, Joshua A. Bauer, Thomas P. Hasaka, Emily L. Days, Reshma R. Desai, Kathryn R. Butcher, Gary A. Sulikowski, Alex G. Waterson, Jens Meiler, Kaitlyn V. Ledwitch, Alfred L. George, Charles R. Sanders

## Abstract

Congenital long QT syndrome (LQTS) promotes risk for life-threatening cardiac arrhythmia and sudden death in children and young adults. Pathogenic variants in the voltage-gated potassium channel KCNQ1 are the most frequently discovered genetic cause. Most LQTS-associated KCNQ1 variants cause loss-of-function secondary to impaired trafficking of the channel to the plasma membrane. There are currently no therapeutic approaches that address this underlying molecular defect. Using a high throughput screening paradigm, we identified VU0494372, a small molecule that increases total and cell surface expression, and trafficking efficiency of wildtype (WT) KCNQ1 as well as three LQTS-associated variants. Additionally, 16-hour treatment of cells with VU0494372 increased current density for WT KCNQ1 and the LQTS-associated variant V207M. VU0494372 had no impact on KCNQ1 transcription, degradation, or thermal stability. We identified a potential direct interaction site with KCNQ1 at or near the binding site of the KCNQ1 potentiator ML277. Together, these findings demonstrate that small molecules can increase the expression levels and cell surface trafficking of KCNQ1 and introduce a potential new pharmacological approach for treating LQTS.

## Introduction

Long QT syndrome (LQTS) is a life-threatening disorder of heart rhythm caused by impaired repolarization of cardiac action potentials, reflected by an increase in duration of the QT interval in the electrocardiogram (1). LQTS is associated with syncope, cardiac arrhythmia, and cardiac arrest, which can be fatal (2). It is estimated that 1:2000-1:2500 individuals are affected by congenital LQTS, making it one of the most common monogenic disorders (3, 4).

Congenital LQTS is caused by pathogenic variants in many genes mostly encoding ion channels or ion channel modulators (5). Type 1 long QT syndrome (LQT1) accounts for up to 50% of cases of congenital LQTS and is caused by loss-of-function of the voltage-gated potassium channel KCNQ1 (also known as K_V_7.1 or K_V_LQT1) (5, 6). KCNQ1 mediates the slow delayed rectifier current (I_Ks_) that drives repolarization during the cardiac action potential (7). Hundreds of pathogenic KCNQ1 variants have been identified, and these variants are distributed throughout the protein sequence (8, 9). Clinical interventions for LQTS include avoidance of triggers such as strenuous exercise, administration of beta blockers, and surgical implantation of a cardioverter-defibrillator (10, 11). However, no treatments currently exist that target the underlying molecular defects in KCNQ1.

Previous studies have indicated that roughly half of known disease-associated variants disrupt KCNQ1 function through protein destabilization, which leads to failure to traffic to the plasma membrane (mistrafficking), in some cases followed by protein degradation (12, 13). In this regard, LQT1 resembles other disorders for which mistrafficking of a membrane protein drives disease mechanisms, including retinitis pigmentosa (rhodopsin photoreceptor) (14–18), diabetes insipidus (vasopressin 2 receptor) (19, 20), type 1E Charcot-Marie-Tooth disease (peripheral myelin protein 22, PMP22) (21–23), and cystic fibrosis (cystic fibrosis transmembrane regulator, CFTR) (24–27). For these proteins, small molecules have been identified that bind to disease-associated variants to stabilize folding and increase cell surface levels (28–34). For example, in cystic fibrosis (CF), extensive research led to the development of a three-drug therapy effective against the ΔF508 CFTR variant responsible for at least 90% of all CF cases (35–37). Two of these three drugs act by rescuing normal folding and trafficking of the mutant protein (trafficking correctors), while the third enhances channel activity (potentiator) (32, 36–38). This treatment has now also been found to be effective for additional CFTR variants (39, 40). As KCNQ1 variants appear to exhibit similar defects in folding and trafficking, we hypothesized that KCNQ1 dysfunction could be remedied in a manner similar to CFTR. While activators of KCNQ1 have been discovered (41–44), no compounds have yet been identified that modify KCNQ1 trafficking. Identification of a small molecule in this class would provide support for this hypothesis and provide proof-of-concept support for treating LQT1 by targeting KCNQ1 dysfunction.

Here, we investigated whether cell surface levels of KCNQ1 and KCNQ1 trafficking efficiency can be pharmacologically enhanced using a high-throughput screen of chemically diverse small molecules. We identified the compound VU0494372, which increases KCNQ1 cell surface levels, total levels, and surface trafficking efficiency in a dose-dependent manner. This compound represents a first-in-class small molecule modulator of KCNQ1 trafficking and provides a foundation for future LQT1 drug discovery efforts.

## Results

### Discovery of KCNQ1 trafficking modulators with high-throughput screening (HTS)

To conduct HTS for compounds that alter the expression and/or trafficking of KCNQ1 in mammalian cells, we developed a high-content image-based trafficking assay. We first generated a cell line that stably expresses KCNQ1 by utilizing “LLP-int” HEK 293T cells, which contain a genomic “landing pad” DNA integration site under control of a tetracycline-inducible promoter where DNA of interest can be integrated (45) (Supplemental Fig. 1A). We inserted a cassette containing wild type human mycKCNQ1-mEGFP cDNA, a KCNQ1 construct with a myc epitope inserted into the extracellular S1-S2 linker (46) and a monomeric, enhanced green fluorescent protein (mEGFP) fused to the C-terminus (Fig. 1A). The cassette also contained an mCherry reporter and a puromycin resistance gene for identification and selection of successfully integrated cells (Supplemental Fig. 1A). The mEGFP fusion allows for visualization and quantification of total KCNQ1 in the cell, while the extracellular myc epitope can be used to visualize/quantify the population of cell surface KCNQ1 using anti-myc antibodies in non-permeabilized cells. Both the mEGFP fusion and the myc epitope insertion have been employed by other laboratories and have been shown not to impact KCNQ1 structure, function, or trafficking (46, 47). We generated a clonal stable cell line by integrating the cDNA for this construct, selecting for successfully integrated cells with puromycin antibiotic selection, sorting single cells into multi-well plates, and growing up a clonal population from a single cell. Inducible expression of mycKCNQ1-mEGFP was confirmed by western blot analysis (Supplemental Fig. 1B).

**Figure 1.**
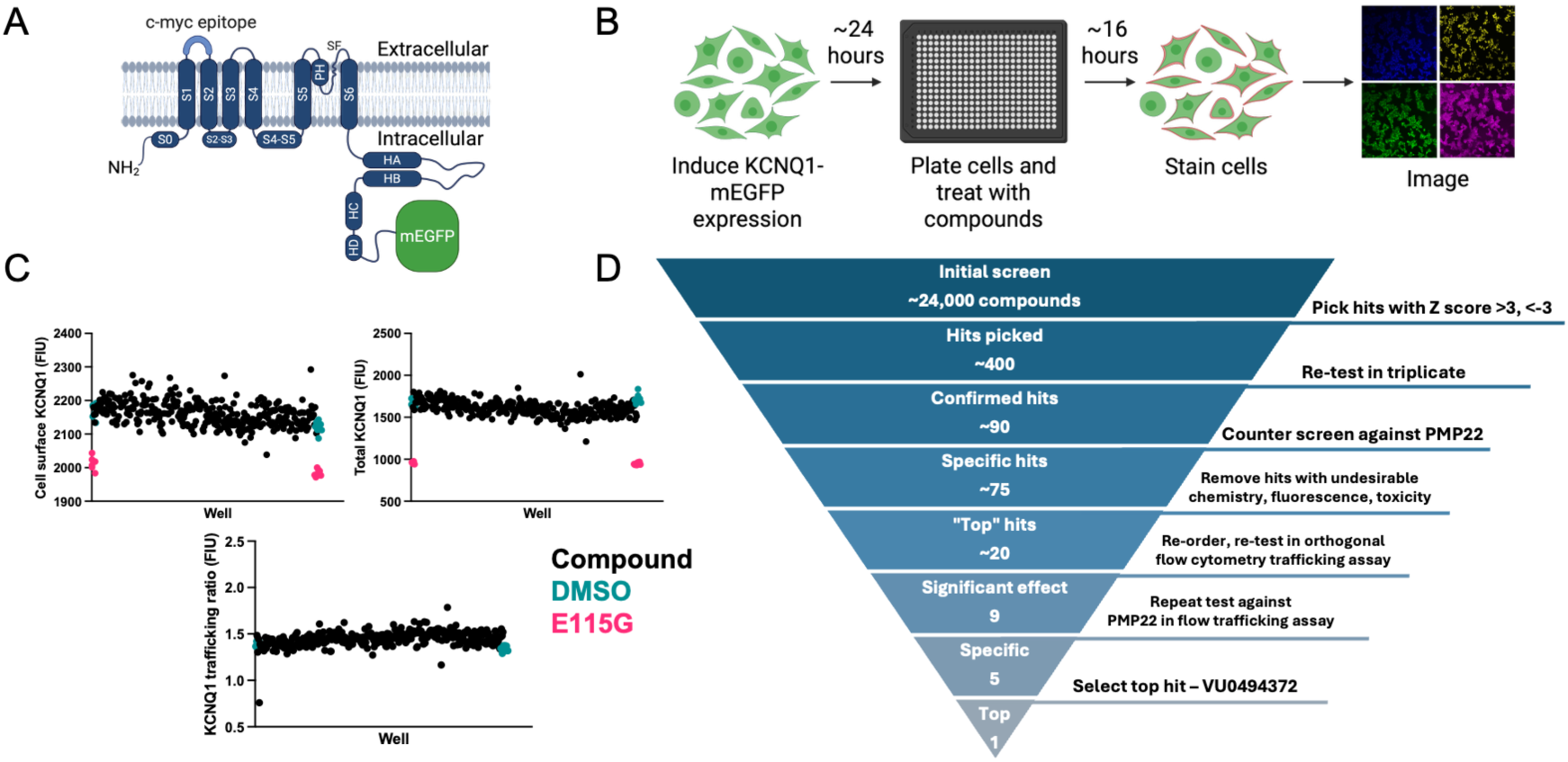
A high throughput screen to identify compounds that alter KCNQ1 expression and/or trafficking. **A)** KCNQ1 construct (mycKCNQ1-mEGFP) used for high-throughput screening. **B)** Workflow for high throughput screening. **C)** Representative data showing quantifications of cell surface KCNQ1 (left), total cellular KCNQ1 (right) and KCNQ1 trafficking ratio (bottom) for a screened 384-well plate. Each point represents the average value for all quantified cells in one well. Black = compound-treated well. Teal = DMSO-treated negative control well. Magenta = E115G mycKCNQ1-mEGFP positive control well. E115G is not shown in the trafficking ratio plot due to skewing of values from near background/level surface and total levels. **D)** Screening funnel summarizing the steps of the process for narrowing from ∼23,000 screened compounds to the top hit VU0494372.

We then used this stable cell line in an immunofluorescence-based trafficking assay compatible with high-content imaging of multi-well plates (Fig. 1B). Expression of mycKCNQ1-mEGFP was induced 24 hours before cells were plated into 384-well plates. Each well was then treated with a screening compound (dissolved in DMSO) or DMSO (control) at a final concentration of 10 µM or 0.1%, respectively. After 16 hours of treatment, cells were fixed, stained, and imaged with a high content imager. Cell surface KCNQ1, total KCNQ1, and the KCNQ1 trafficking ratio (surface fluorescence intensity/total fluorescence intensity) were quantified for each imaged cell, and an average surface, total, and trafficking ratio value was calculated for each compound. Representative data from a screened plate is shown in Figure 1C.

To validate the assay, a clonal stable cell line expressing E115G mycKCNQ1-mEGFP, a LQT1-associated variant with known low total and cell surface levels of KCNQ1 (13), was used as a positive control. Consistent separation between negative control (DMSO-treated WT) cells and the positive control was observed (Supplemental Fig. 1C). Cells were also tested to ensure that treatment with 0.1% DMSO did not alter mycKCNQ1-mEGFP surface, total, or efficiency values prior to screening (Supplemental Fig. 1D).

A total of 23,611 small molecules were screened from five libraries. Robust Z scores were calculated for cell surface KCNQ1, total KCNQ1, and KCNQ1 surface trafficking ratio for each screened compound on a plate-by-plate basis (Supplemental Fig. 2). Compounds with a robust Z score >3 or <-3 for any of the three quantified metrics (surface, total, trafficking ratio) that were not toxic were selected as initial hits (n=386, 1.6% hit rate). A series of experiments were then performed to narrow the list of hits (Fig. 1D). Hits were re-tested in triplicate, and 87 compounds with reproducible effects were identified. These hits were then counter-screened against an unrelated membrane protein, PMP22, to identify those that also altered PMP22 expression/trafficking, indicating non-specific effects. Such non-specific hits were excluded from further studies. Finally, for the top 21 hit compounds that met all desired criteria, a fresh supply of material was obtained from commercial sources. The new material was tested in a flow cytometry-based trafficking assay, which allowed for more precise quantification of protein levels and cell surface expression. Five compounds were found to have reproducible, specific effects (Supplemental Fig. 3). Three of the top hits decreased cell surface and total KCNQ1 levels, and another increased total KCNQ1 dramatically but led only to a small increase in cell surface levels, leading to a net decrease in trafficking efficiency. The fifth hit, VU0494372 (Fig. 2), increased cell surface and total KCNQ1 levels as well as KCNQ1 trafficking efficiency. It is this hit that we focus on in this paper.

**Figure 2.**
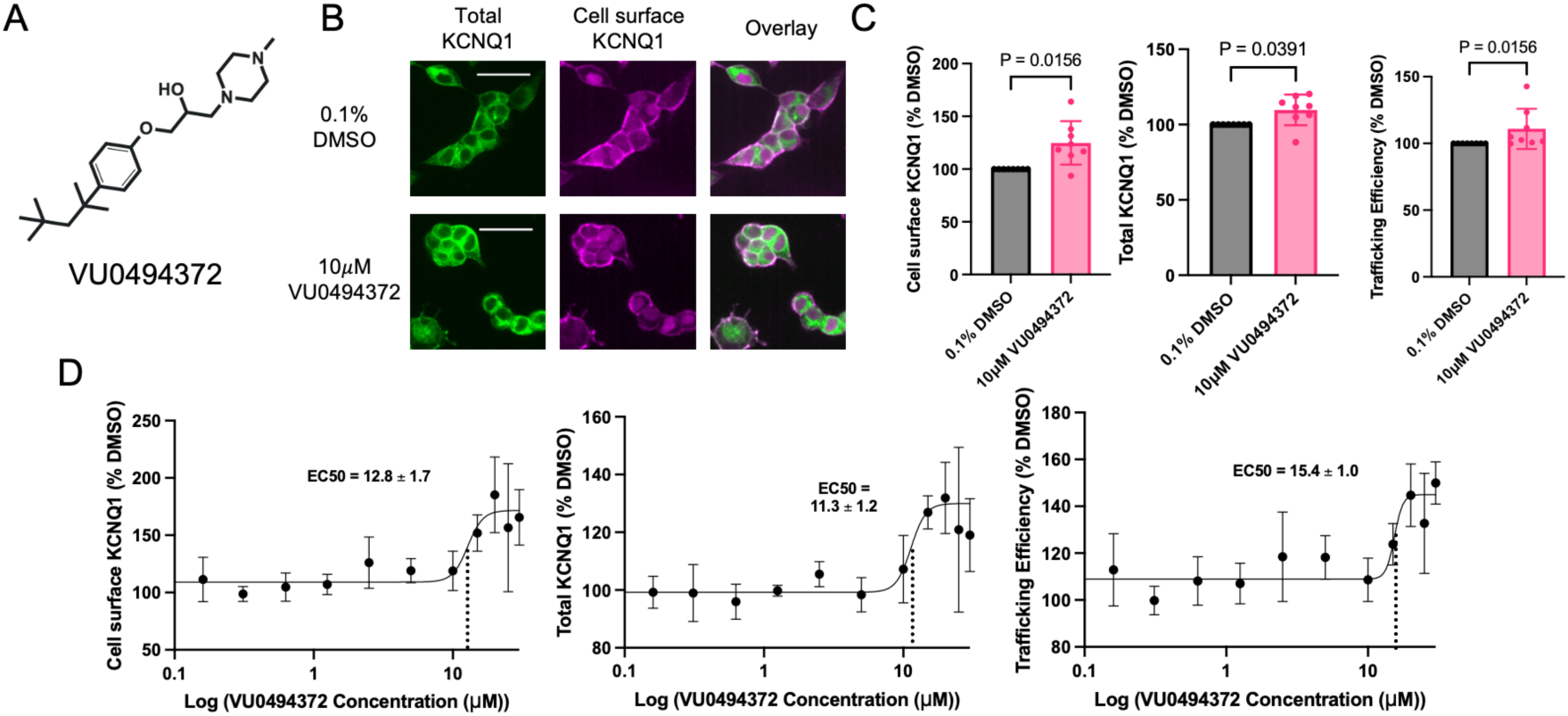
VU0494372 increases cell surface and total KCNQ1 in a dose-dependent fashion. **A)** Chemical structure of VU0494372. **B)** Images of total (left) or cell surface (middle) mycKCNQ1-mEGFP in LLP-int cells treated with either 0.1% DMSO (top) as a control or 10 µM VU0494372 (bottom). Scale bar represents 40 µm and scale is the same in all images **C)** Expression and trafficking of mycKCNQ1 in stable cell line, as determined by flow cytometry-based trafficking assay. Values were normalized to 0.1% DMSO control within each replicate. P values were determined with Wilcoxon matched-pairs signed rank test. N=8. Error bars represent standard deviation. **D)** Dose-response curves for mycKCNQ1 cell surface levels (left), total levels (middle), and trafficking efficiency (right), in VU0494372-treated stable cell line. Values were normalized to 0.2% DMSO control in each replicate. The best-fit curve was generated and EC_50_ values were determined with GraphPad Prism. The curve in the “total” plot was constrained to have a maximum of 130%, and the curve in the “trafficking” plot was constrained to have a maximum of 145% due to large variability at high concentrations. N=3-5. Error bars represent standard deviation.

### VU0494372 increases cell surface KCNQ1, total KCNQ1, and KCNQ1 trafficking efficiency in a dose-dependent manner

We first verified the effect of VU0494372 in the quantitative flow cytometry-based trafficking assay using a stable cell line expressing WT mycKCNQ1 (without mEGFP) to ensure the effect of VU0494372 was not due to an interaction with mEGFP. We found that 10 µM VU0494372 treatment led to a 25% increase in cell surface KCNQ1, a 9.7% increase in total mycKCNQ1, and an 11% increase in mycKCNQ1 trafficking efficiency (Fig. 2C). VU0494372 did not impact the unrelated membrane protein PMP22, indicating that the compound is not having a pleiotropic effect on membrane protein trafficking (Supplemental Fig. 4). We also determined that VU0494372 exhibited a dose-dependent effect with a half maximal effective concentration (EC_50_) of 12.8 ± 1.7 µM for increased cell surface KCNQ1 levels, an EC_50_ of 11.3 ± 1.2 µM for increased total KCNQ1 levels, and an EC_50_ of 15.4 ± 1.0 µM for increased KCNQ1 trafficking efficiency (Fig. 2D). Together, these results demonstrate that VU0494372 increases cell surface levels, total levels, and trafficking efficiency of WT KCNQ1 in a dose-dependent manner. We next sought to determine whether VU0494372 was also effective for disease-associated KCNQ1 variants.

### Treatment with VU0494372 increases cell surface levels and trafficking efficiency of LQT1-associated variants

We next determined the effect of VU0494372 on the surface levels, total levels, and trafficking efficiency of LQT1-associated KCNQ1 variants. We selected three previously characterized variants: G179S and G189E, known to be severely dysfunctional in trafficking and function, and V207M, known to have a milder, but still disease-causing, phenotype (12, 13). We generated “population” stable cell lines with WT mycKCNQ1 and the three LQT1-associated variants similarly to our approach for the original stable cell lines but omitting the final clonal population growth step. Cells were treated with 20 µM VU0494372 (a concentration above the determined EC_50_, to ensure maximal effect size) for 16 hours. We found that treatment with 20 µM VU0494372 resulted in significantly increased cell surface KCNQ1 and KCNQ1 trafficking efficiency for all three LQT1 variants, and no change in total KCNQ1 (Fig. 3A-C). The variants exhibited a greater response to VU0494372 compared to the 1.5-fold increase for WT KCNQ1. G189E and G179S exhibited >2-fold and >3-fold increases in cell surface levels, respectively, while V207M exhibited an approximately 2-fold increase. Increases in trafficking efficiency represented a similar trend, with all three variants exhibiting a larger magnitude change in trafficking than WT after treatment with VU0494372. These results demonstrate that these three LQT1-associated KCNQ1 variants are responsive to VU0494372 treatment and exhibit even larger increases in surface and trafficking values than WT. After determining the effects of VU0494372 on WT and variant KCNQ1, we investigated its mechanism of action.

**Figure 3:**
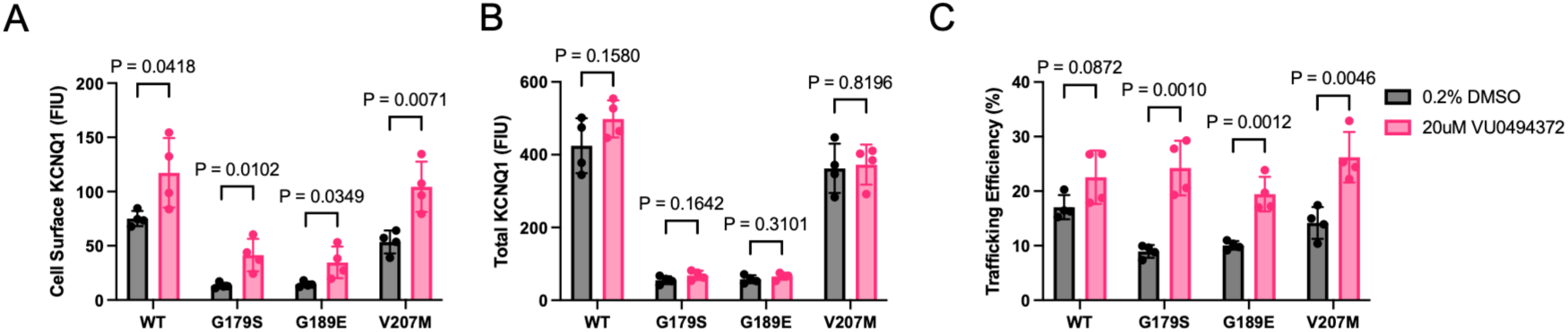
Treatment with VU0494372 increases cell surface levels and trafficking efficiency of LQT1-associated variants. Cell surface **(A)**, total **(B)**, and trafficking efficiency **(C)** data for WT, G179S, G189E, and V207M mycKCNQ1 in “population” stable cell lines, as determined by flow cytometry-based trafficking assays, after treatment with 0.2% DMSO or 20 µM VU0494372 for 16 hours. N=4. Error bars represent standard deviation, and P values were determined with unpaired t-tests.

### VU0494372 does not alter KCNQ1 transcription, degradation, or stability

To investigate the mechanism of action, we conducted a series of studies testing the impact of VU0494372 on KCNQ1 expression, degradation, and stability. Because treatment with VU0494372 increased total levels of KCNQ1, we investigated whether this change was due to increased KCNQ1 production or a reduction in KCNQ1 degradation. We began by examining whether the increased levels of KCNQ1 in cells were a result of VU0494372 altering KCNQ1 at the transcriptional level. We used RT-qPCR to quantify KCNQ1 mRNA levels in our stable cell line and found that treatment with VU0494372 did not alter the levels of KCNQ1 mRNA (Fig. 4A). We next performed cycloheximide chase assays, which involve inhibiting synthesis of new protein and tracking target protein degradation over time, to determine the effect of VU0494372 on the rate of degradation of KCNQ1. These experiments determined that the cellular half-life of WT KCNQ1 was approximately 7 hours, and that treatment with VU0494372 did not change the rate of KCNQ1 degradation (Fig. 4B).

**Figure 4:**
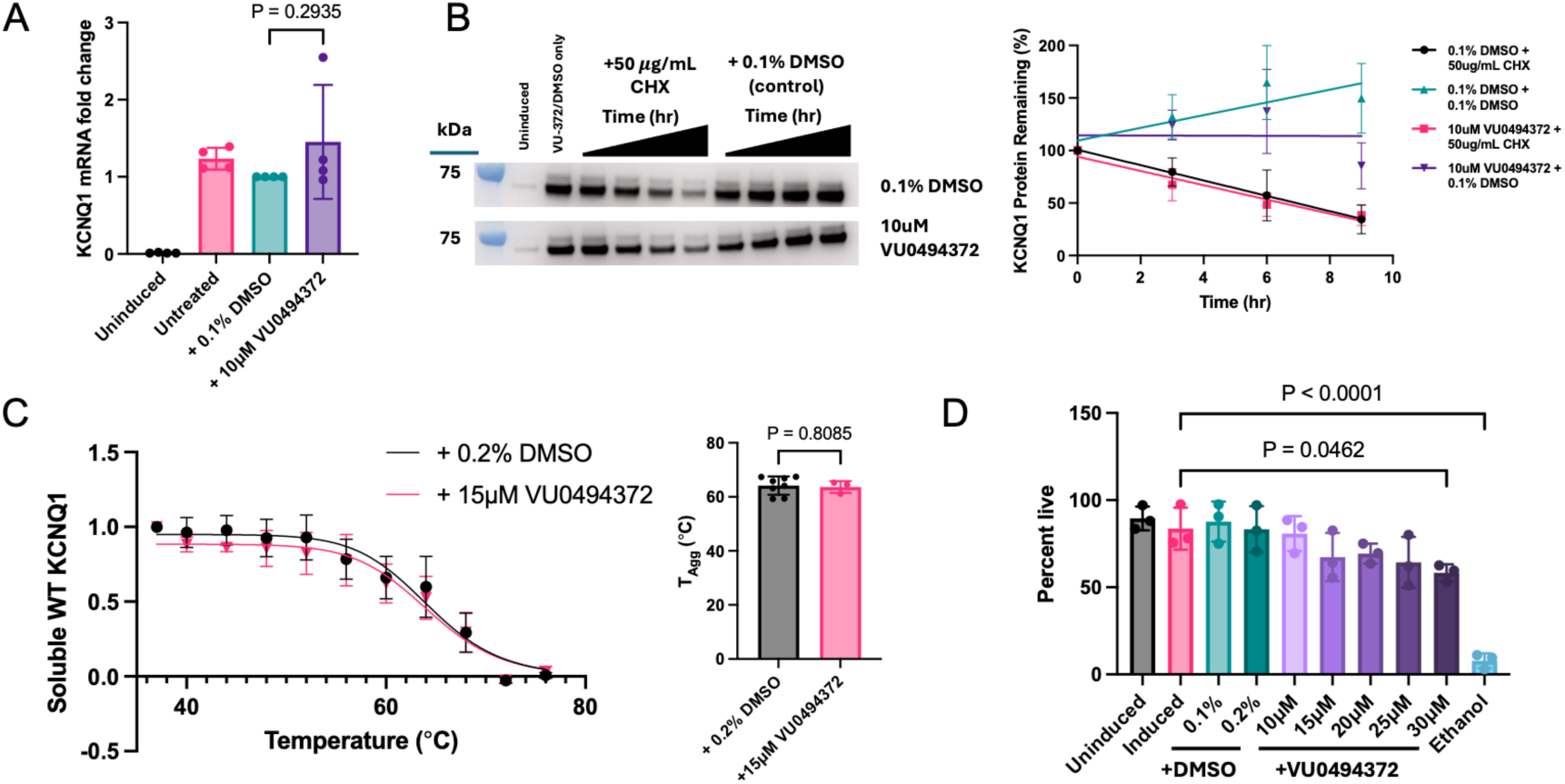
VU0494372 effects do not involve modulation of KCNQ1 degradation, transcription, thermal stability, or cell toxicity. **A)** Fold-change in KCNQ1 mRNA levels, as determined by qPCR, relative to 0.1% DMSO-treated samples. N=4. P values were calculated with Kruskal-Wallis tests with Dunn’s multiple comparisons test as follow-up. Error bars represent standard deviation. **B)** Cycloheximide chase assay to determine the rate of KCNQ1 degradation. Remaining KCNQ1 was detected with western blotting (left). Representative full blots are shown in Sup. Fig. 5. KCNQ1 band intensities were normalized to beta-actin loading control bands and plotted as a percent remaining from the 0-hour time point (right). N=3 replicates, for which error bars represent standard deviation. Lines of best fit were generated with GraphPad Prism. **C)** Thermal aggregation curve (left) from cellular thermal shift assay (CETSA) experiments. Bands corresponding to KCNQ1 were quantified and T_Agg_ values were calculated for N=3 (VU0494372-treated) or 5 (DMSO-treated) replicates. Curve of best fit modeled with GraphPad Prism. T_Agg_ quantifications (right) were calculated based on the inflection point of the modeled curves in each individual CETSA replicate. An unpaired t-test was used to determine the P value. Error bars on both graphs represent standard deviations. Full representative blots are shown in Sup. Fig. 5 **D)** Toxicity of 0.1% - 0.2% DMSO or 10 – 30 µM VU0494372, determined by using trypan blue to assess cell viability. One-way ANOVA with Dunnett’s multiple comparisons follow-up test was used to determine P values. N=3. Error bars represent standard deviation.

Because the overall efficiency of trafficking of KCNQ1 was increased, we sought to determine if this was mediated by an alteration in the stability of KCNQ1, which could facilitate proper folding and trafficking. A cellular thermal shift assay (CETSA) was utilized to calculate a thermal aggregation temperature (T_Agg_) for KCNQ1 under cellular conditions with and without 15 µM VU0494372. Treatment with VU0494372 did not alter the thermal stability of KCNQ1 (Fig. 4C).

Finally, to rule out toxicity-based mechanisms (e.g., apoptosis) as a confounding factor in the effect of VU0494372, we tested whether VU0494372 induced significant cellular toxicity. Cells were treated with 10 – 30 µM VU0494372 for 16 hours followed by assessment of viability using trypan blue staining. The compound did not significantly impact the viability of the mycKCNQ1-expressing stable cell line at concentrations of up to 25 µM (Fig. 4D).

Together, these experiments rule out alterations in transcription, degradation, or global stability as possible mechanisms of action for VU0494372 and show that VU0494372 is not significantly toxic to cells at the concentrations used. We next investigated whether VU0494372 impacted the function of KCNQ1, which could provide further insight into its mechanism of action.

### Functional effects of acute and chronic VU0494372 exposure

After determining that VU0494372 increases cell surface levels of WT and LQT1-associated KCNQ1 variants, we investigated whether VU0494372 treatment led to corresponding changes in KCNQ1 I_Ks_ currents. Automated whole-cell patch clamp electrophysiology experiments were conducted with CHO cells stably expressing KCNE1 (CHO-KCNE1) and transiently transfected with WT KCNQ1. Cells were treated with 20 µM VU0494372 for 16 hours (chronic treatment) to ensure maximal effect size and allow time for alterations in cell surface levels to occur. VU0494372 was removed before recording currents. We found that WT KCNQ1 peak current density significantly increased after treatment with VU0494372, and the V_1/2_ of activation of KCNQ1 shifted significantly to less depolarized potentials after treatment (Fig. 5A). These effects were dose-dependent, although full dose-response curves could not be obtained due to toxicity at concentrations greater than 20 µM (Sup. Fig. 6). The increase in peak current density likely reflects an increase in the number of channels at the cell surface, and the shift in the V_1/2_ of activation of KCNQ1 reflects a change in the gating of KCNQ1, which could be occurring through any one of several possible mechanisms (see Discussion).

**Figure 5:**
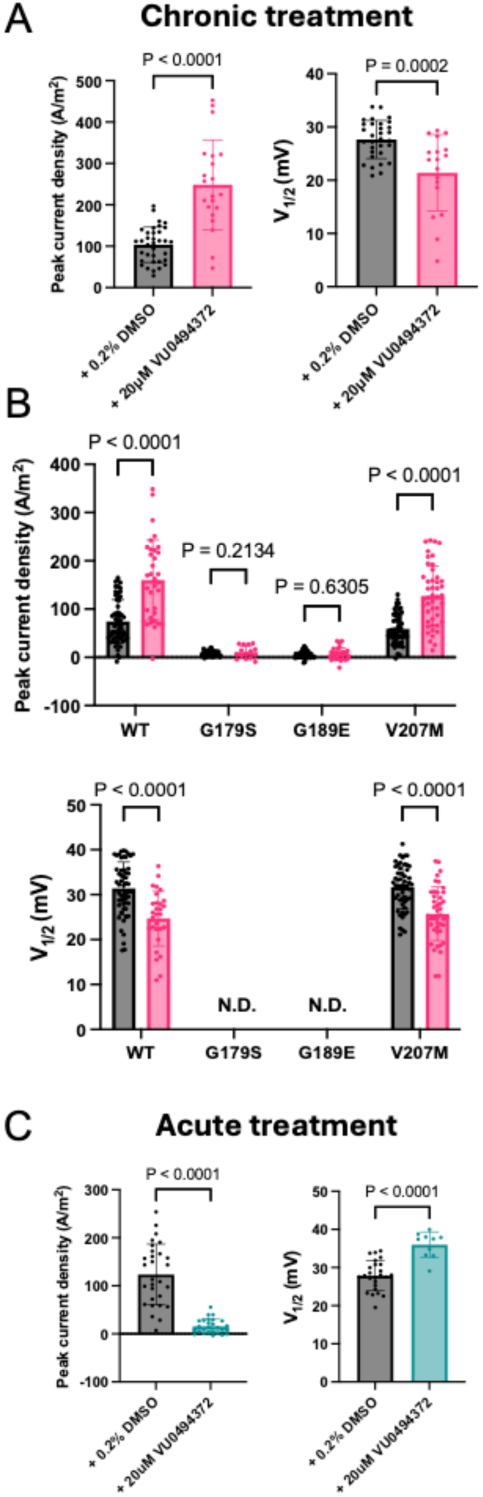
Treatment with VU0494372 leads to an increase in WT and V207M KCNQ1 activity. A-C) KCNQ1 channel function quantified in CHO-K1 cells stably expressing KCNE1 (CHO-KCNE1) and transfected with WT or variant KCNQ1 using automated patch clamp electrophysiology (n=10-60 cells per group). Error bars represent standard deviation and P values were determined with unpaired t-tests. **A)** Peak current density (left) and V_1/2_ of activation (right) of WT KCNQ1 after 16-hour (chronic) treatment with 0.2% DMSO or 20 µM VU0494372. Compound was removed before recording (n=18-35 cells were quantified from 2 separate experimental replicates). **B)** Peak channel current density (top) and V_1/2_ of activation (bottom) for CHO-KCNE1 cells transiently transfected with WT or LQT1 variant G179S, G189E, or V207M KCNQ1 after chronic treatment with 0.2% DMSO or 20 µM VU0494372. Compound or DMSO was washed off before recording (n=15-60 cells per group). N.D.= not determined, due to low current amplitude. **C)** Peak current density (left) and V_1/2_ of activation (right), as determined for CHO-KCNE1 cells transfected with WT KCNQ1. Cells were treated with 0.2% DMSO or 20 µM VU0494372 for 5 minutes (acute treatment) before recording, and treatment remained present during recording (n=10-29).

We next determined whether VU0494372 alters the function of disease-associated variants of KCNQ1. CHO-KCNE1 cells transiently transfected with WT, G179S, G189E, or V207M KCNQ1 were treated with 20 µM VU0494372 for 16 hours, followed by removal of the compound. Automated whole-cell patch clamp experiments demonstrated that chronic treatment of V207M KCNQ1 led to significant increases in current and depolarization of activation V_1/2_ similarly to WT. However, G179S and G189E did not exhibit changes in KCNQ1 current density, and their V_1/2_ of activation could not be determined due to low overall current (Fig. 5B). This suggests that even though surface trafficking of these variants is very significantly enhanced, the surface-trafficked population of the channel remains dysfunctional.

Automated whole-cell patch clamp studies of CHO-KCNE1 cells transiently transfected with WT KCNQ1 and treated with 0.2% DMSO or 20 µM VU0494372 for 5 minutes (acute exposure, no washing) were also conducted. For this experiment, VU0494372 or DMSO (control) was added 5 minutes prior to whole-cell recordings and maintained throughout the experiment. These experiments showed the opposite effect of chronic treatment - a decrease in KCNQ1 peak current density and a positive shift in V_1/2_ of activation (Fig. 5C). These changes were consistent with direct inhibition by VU0494372 of KCNQ1 channel function. The decrease in KCNQ1 currents after this short-term exposure strongly suggests that the small molecule is interacting directly with KCNQ1 to reduce channel activity, while the shift in the activation V_1/2_ reflects alteration of the voltage sensitivity of gating.

Together, our electrophysiological studies demonstrate that chronic treatment with VU0494372, followed by compound removal, leads to increases in WT and V207M KCNQ1 function, consistent with increased trafficking to the plasma membrane. On the other hand, acute continuous exposure leads to decreases in WT KCNQ1 function, indicating that VU0494372 can act as a channel blocker while increasing trafficking efficiency. This led us to next investigate whether VU0494372 is directly interacting with KCNQ1.

### Competition assays support direct interaction between VU0494372 and KCNQ1

The observation that acute, continuous exposure of KCNQ1-expressing CHO-KCNE1 cells to VU0494372 inhibits channel function suggests that this compound directly interacts with KCNQ1. To strengthen evidence for direct interaction, we used a combination of experimental and computational molecular modeling approaches to predict and evaluate potential binding sites of VU0494372 to KCNQ1. VU0494372 was blind-docked to two different cryo-EM structures of KCNQ1, representing two different conformations of the protein; a voltage sensor up with pore closed conformation (PDB 8SIK) (48), and a voltage sensor active state with pore open conformation (PDB 7XNK) (49). We used DiffDock (50) to generate predicted binding poses. We then refined and scored the DiffDock poses with the RosettaLigand (51, 52) small molecule docking protocol for high-resolution refinement. The best scoring poses with the most favorable binding energies positioned VU0494372 proximal to the KCNQ1 pore domain, forming contacts with the S6 transmembrane helix and the S5-S6 linker. (Fig. 6, A-C).

**Figure 6:**
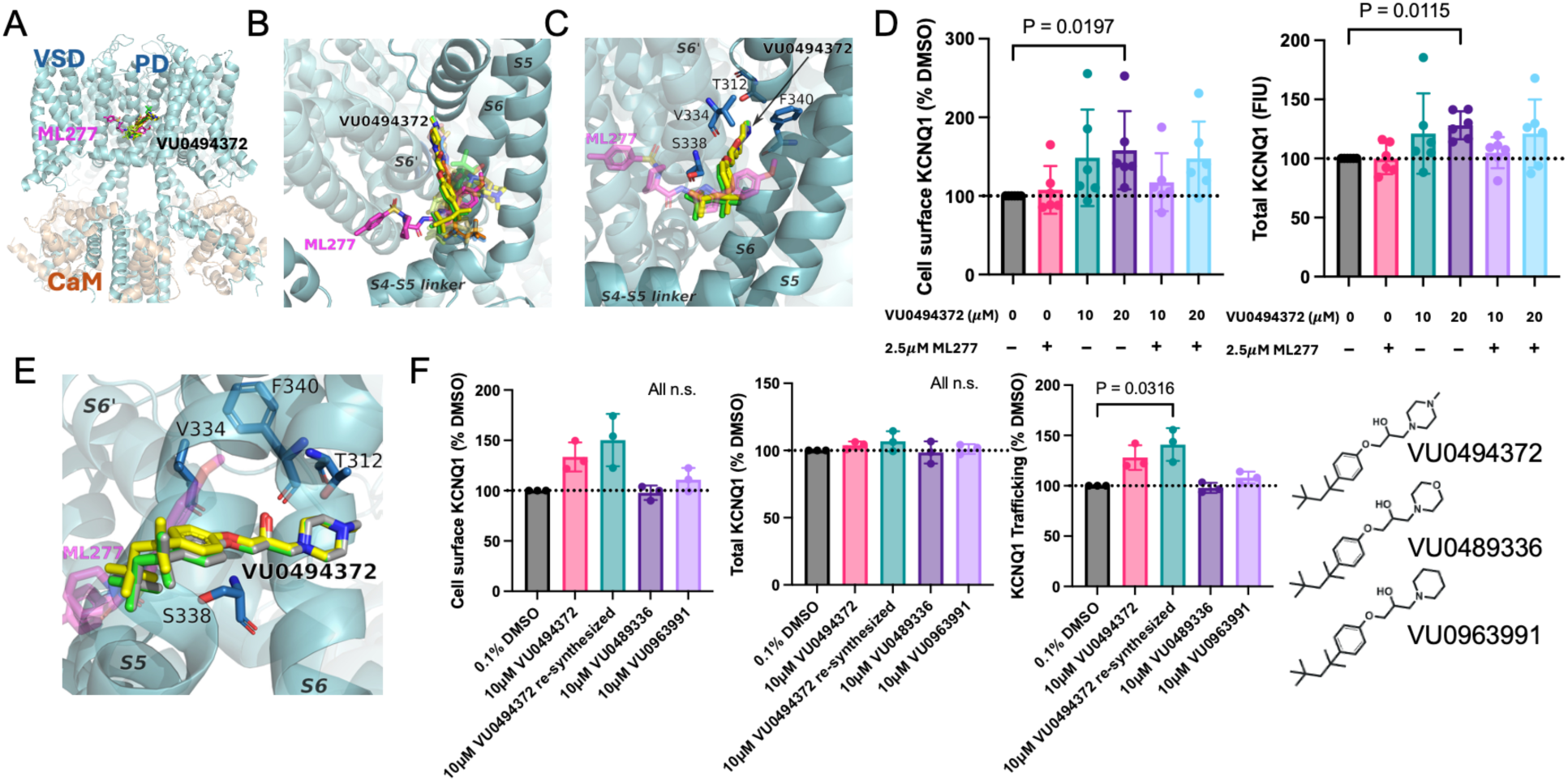
Computational docking, electrophysiology data, and competition assays provide support for possible direct interaction between VU0494372 and KCNQ1. **A)** Overlay of PDB 7XNK showing ML277 (magenta) bound to voltage sensor active, pore open KCNQ1 and the top ten docked poses of VU0494372 (assorted colors) ranked by RosettaLigand interface score. VSD: Voltage-sensing domain. PD: pore domain. CaM: calmodulin. **B)** Zoomed and rotated view of panel A. Docked VU0494372 poses overlap with bound ML277 in KCNQ1. Labeled residues line the predicted pocket and interact with VU0494372. **C)** Four poses from panel B of the most frequently observed orientation of VU0494372 docked to KCNQ1. **D)** Competition between ML277 and VU0494372 as determined with a flow cytometry-based trafficking assay. Cell surface (left) and total (right) mycKCNQ1 levels in LLP-int cells were determined with and without ML277 and VU0494372. N=6. P values were determined with Kruskal-Wallis nonparametric test with Dunn’s multiple comparison test for follow-up. **E)** Rotated and zoomed view of panel C to illustrate the orientation of VU0494372. **F)** Cell surface levels (left), total levels (middle), and trafficking efficiency (right) of WT KCNQ1 in a LLP-int WT KCNQ1-expressing cell line after treatment with 10 µM VU0494372, VU0489336, or VU0963991 (or 0.1% DMSO as a control) for 16 hours, quantified with a flow cytometry-based trafficking assay. N=3. Compound structures are shown on the far right. P values were determined with Kruskal-Wallis nonparametric test with Dunn’s multiple comparison test for follow-up.

All docked poses partially overlapped with the binding site of a known KCNQ1 activator, ML277 (41) (Fig. 6, A-C). ML277 has an EC_50_ of 0.26 µM and does not alter the trafficking of KCNQ1, even at concentrations as high as 10x the EC_50_ (Fig. 6D) (41). We therefore investigated whether ML227 and VU0494372 compete for the same binding site. We treated LLP-int mycKCNQ1 cells with 2.5 µM ML277 and either 10 µM or 20 µM VU0494372 and used our flow cytometry trafficking assay to quantify cell surface and total KCNQ1. We then compared these values to those from ML277 or VU0494372 alone and found that co-treating with ML277 reduced the effects of VU0494372 on KCNQ1 levels (Fig. 6D), consistent with the notion that the two small molecules are competing for the same binding site.

We also sought to determine whether alterations to the chemical substituents of VU0494372 predicted to interact with KCNQ1 impacted its effect on the channel, which could indicate altered binding. One end of the compound, containing a methylpiperazine ring, was predicted to penetrate into KCNQ1 and interact with several residues in the KCNQ1 pore domain (Fig. 6E). The Vanderbilt Molecular Design and Synthesis Core synthesized VU0494372 and two analogs (VU0489336 and VU0963991) with modifications to this methylpiperazine ring present in the parent compound (Fig. 6F). These two analogs exhibited reduced effects on cell surface KCNQ1 and trafficking efficiency (Fig. 6F), suggesting that this portion of the compound was essential for maximal effect of VU0494372 on surface trafficking, possibly due to direct interaction with KCNQ1 residues in the pore domain.

Combined, these data suggest that VU0494372 interacts directly with KCNQ1 near the ML277 binding site with the methylpiperazine ring oriented toward the channel pore, potentially interacting with residues including T312, V334, and F340 in the S6 helix.

## Discussion

### A versatile HTS discovery assay for compounds that alter KCNQ1 levels and/or trafficking

In this work, we used a high content image-based HTS trafficking assay to test the hypothesis that KCNQ1 expression and/or trafficking can be altered with small molecules, a result that provides proof-of-concept support for the potential use of small molecules of this class to remedy KCNQ1 dysfunction in LQT1. We screened nearly 24,000 small molecules for their impact on WT KCNQ1 cell surface levels, total levels, and trafficking efficiency. This approach is advantageous because it enables identification of compounds that target a single aspect of KCNQ1 expression and trafficking, such as changing surface levels without altering total levels. This assay is additionally useful because it is bidirectional; it can be used to identify small molecules that increase KCNQ1 levels and trafficking and are relevant to LQTS as well as small molecules that lower KCNQ1 levels and/or trafficking efficiency. In fact, three of the top five hit compounds decreased surface and total KCNQ1 levels (Supplemental Fig. 3). Compounds of this latter class are of interest as potential drug leads for rare cardiac disorders such as short QT syndrome and familial atrial fibrillation, where the disease is caused by excessive KCNQ1 expression and/or trafficking or by unregulated KCNQ1 channel function (53, 54). This assay may also be adapted for use with more complex systems such as cardiomyocytes or organoids.

### VU0494372 enhances both total and surface levels of KCNQ1 and increases its surface trafficking efficiency

VU0494372 is a small molecule identified by our HTS screen that, at 20 µM (near 2X EC_50_), increases WT KCNQ1 cell surface levels by about 40% (Fig. 2 and 3). Moreover, treatment with 20 µM VU0494372 for 16 hours led to enhancements of 2-fold and higher for cell surface levels and trafficking efficiency for the disease-linked variants G179S, G189E, and V207M (Fig. 3). These effect sizes were larger than those observed for WT KCNQ1, suggesting that disease-associated variants in KCNQ1 may be more sensitive to the compound than WT. Additional results ruled out several possible artifactual explanations for how VU0494372 enhances KCNQ1 trafficking and indicate that VU0494372 specifically targets KCNQ1, appearing to bind directly (Fig. 6). Additionally, our studies found that VU0494372 does not change the rate of KCNQ1 degradation nor broadly alter KCNQ1’s thermal stability in cells (Fig. 4).

While a number of KCNQ1 activators and blockers have previously been identified (41–44, 55, 56), VU0494372 is the first small molecule that increases cell surface levels of KCNQ1. These results support our hypothesis that KCNQ1 expression and trafficking can be modified by small molecules and suggest that pharmacological enhancement of KCNQ1 trafficking is a potential route for treating LQT1. It is particularly encouraging both that the studied disease-associated KCNQ1 variants exhibit larger improvements in surface trafficking than WT and also that all three tested variants were responsive to VU0494372. It is estimated that about half of the hundreds of known LQT1-associated KCNQ1 variants exhibit impaired cell surface expression (12, 13), so a small molecule trafficking corrector that remedies these variants would be broadly useful for treating LQT1.

### VU0494372 increases WT KCNQ1 function after chronic treatment

Our functional studies show that treatment of KCNQ1/KCNE1 co-expressing CHO cells with VU0494372 for 16 hours followed by compound removal results in increased KCNQ1 current (Fig. 5). However, a depolarizing shift in the activation V_1/2_ of the I_Ks_ channel was also observed (Fig. 5A). This change in the voltage sensitivity of channel gating could be the result of any one of several mechanisms, including either drug-induced alteration of the phosphorylation of the channel or stabilization of an interaction with a chaperone or accessory protein, both of which have previously been shown to shift the V_1/2_ of activation for KCNQ1 (57–59). Overall, these changes in V_1/2_ also represent a potentiating effect, though the mechanism is not yet clear. A similar effect has been observed for the KCNQ2 activating drug retigabine (60, 61).

Additionally, the effect of this compound in CHO cells co-expressing KCNE1 and KCNQ1 confirms that VU0494372 still exerts its effect on KCNQ1 in the presence of its accessory protein KCNE1, despite initially being identified in HTS screens conducted in the absence of KCNE1. Effectiveness on both KCNQ1 alone and KCNQ1-KCNE1 does not occur for some modulators of KCNQ1. For example, ML277 and R-L3’s effects on KCNQ1 function are hindered in the presence of KCNE1 (43, 62). Since the KCNQ1-KCNE1 complex is the active form of the channel in the heart (59), this result provides additional evidence that small molecules such as VU0494372 have potential for clinical use and benefit.

### Variant-specific dependence of the impact of VU0494372 on channel function

While VU0494372 increased the cell surface levels and trafficking efficiency of three LQT1-associated variants: G179S, G189E, and V207M (Fig. 3), only V207M showed significant increases in channel conductance after treatment with VU0494372 (Fig. 5B), despite cell surface levels and trafficking efficiency improving by 2-fold or more for all three studied variants. This can likely be explained by the degree of dysfunction of each variant, which has been previously characterized (12, 13). G179S and G189E are known to cause severe dysfunction of KCNQ1; we have previously demonstrated that not only are both destabilized and have poor cell surface trafficking, they also exhibit extremely low KCNQ1 conductance (12, 13). By contrast, V207M exhibits a less severe phenotype with only modest impairments in trafficking, function, and stability (13).

Our results suggest that some “milder” disease-associated variants, like V207M, that mistraffic but are still able to function may be susceptible to rescue by compounds that modulate expression/trafficking. Multi-pronged approaches like treating with both a trafficking modulator and a channel activator may be needed to remedy more severely dysfunctional variants such as G179S and G189E. This dual approach has been utilized for variants in CFTR that cause cystic fibrosis, in particular the common ΔF508 form, which cause both mistrafficking and loss of function of the channel (35–37, 63).

### Interaction of VU0494372 with KCNQ1

Our data suggest direct interaction between KCNQ1 and VU0494372 as the basis for mediating its mechanism of action, as functional testing after acute, continuous exposure treatment with VU0494372 demonstrated an inhibitory effect (Fig. 5C). Computational docking of VU0494372 to KCNQ1 identified a favorable binding site between the voltage sensing and pore domains of KCNQ1 with the methylpiperazine ring of VU0494372 making contacts with pore domain residues (Fig. 6A-C,E). Subsequent competition experiments with the KCNQ1 activator ML277 (49, 64), which is known to bind near this site, and examination of VU0494372 analogs with modifications to the methylpiperazine ring that further reduced activity supported the potential for direct interaction (Fig. 6D,F).

It cannot be ruled out that the apparent competitive effects of VU0494372 and ML277 are seen for a reason other than direct competition, such as allostery. Further experiments specifically focusing on the impact of mutations in the proposed binding site and/or equilibrium binding studies will be needed to confirm this direct interaction.

### Limitations of VU0494372

While the ability of VU0494372 to enhance cell surface trafficking of KCNQ1 represents an encouraging development, this compound does exhibit some limitations as a potential lead for small molecule drug discovery. An addressable limitation is that the EC_50_ of the effect of VU0494372 on KCNQ1 trafficking is in the range of 10-15 µM, higher than is desirable for a viable drug candidate. It is possible that the potency of this compound can be improved by iterative medicinal chemistry and analog testing. A modest first step toward these SAR studies is illustrated by data in Fig. 6F.

A more serious limitation is the fact that VU0494372 acts as a channel blocker (Fig. 5C), albeit reversibly so (Fig. 5A). This phenomenon has been previously observed with some of the original “pharmacological chaperones”, which increased surface trafficking of several different G protein-coupled receptors, even though they were known receptor antagonists (33, 65). Small molecule-induced increases in trafficking combined with acute channel block have also been observed for potassium channel hERG (K_V_11.1), variants of which cause type 2 long QT syndrome (5). Several drugs, including astemizole (an antihistamine), dofetilide (an approved anti-arrhythmia drug), and E4031 (an investigational anti-arrhythmia drug), directly bind hERG, increase its trafficking, and also block channel function (66), likely by binding to a site important for channel function or by blocking the channel pore (66). In any case, this KCNQ1 channel inhibitory property would likely need to be eliminated through SAR before a compound derived from VU0494372 could be used to treat LQT1. Nevertheless, identification of VU0494372 as a corrector of KCNQ1 trafficking is encouraging for future LQT1 drug discovery efforts.

## Methods

### Cell culture conditions

All mammalian cells were cultured at 37 °C and 5% CO_2_. HEK 293T “LLP-int” cells, and all stable cell lines generated from LLP-int cells, were cultured in high glucose Dulbecco’s Modified Eagle Medium (DMEDM) without sodium pyruvate (Gibco) supplemented with 100 U/mL penicillin/streptomycin (Gibco) and 10% tetracycline negative FBS (Corning) and grown in flasks coated with poly-L-lysine (PLL, Millipore). HEK 293 cells were cultured in high glucose DMEM without sodium pyruvate (Gibco) with pen./strep. And FBS. CHO-K1 cells constitutively expressing human KCNE1 (designated CHO-KCNE1 cells) used for electrophysiology experiments were generated as previously described (67) using the FLP-in^TM^ system (Thermo Fisher Scientific) and were grown in F-12 Ham nutrient medium (Gibco/Invitrogen) supplemented with 10% FBS (ATLANTA Biologicals), penicillin (50 units/mL), streptomycin (50 µg/mL) and maintained under selection with hygromycin B (600 µg/mL).

### KCNQ1 constructs and mutagenesis

All experiments were conducted with “mycKCNQ1”: full-length human KCNQ1 (GenBank accession number AF000571) with a myc epitope (EQKLISEEDL) inserted after residue E146 in the S1-S2 extracellular loop of the KCNQ1 voltage sensing domain (46), or “mycKCNQ1-mEGFP”: mycKCNQ1 with a monomeric enhanced green fluorescent protein (mEGFP, containing the A206K mutation) fused to the C-terminus. Fusion protein generation is described in detail in the Supplemental Information.

### Landing pad cells

Stable cell lines expressing mycKCNQ1-mEGFP WT, mycKCNQ1-mEGFP E115G, and mycKCNQ1 WT, and G179S, G189E, and V207M mycKCNQ1 used for high-throughput screening (HTS) and subsequent experiments were generated using “LLP-int” HEK 293T cells, described in detail in previous publications (45, 68), which contain a tetracycline-inducible genomic “landing pad” DNA integration site and a Bxb1 integrase gene. Target DNA flanked by a complementary integration sequence in a promoterless “shuttle vector” can be incorporated at this site. Details of stable cell line generation are described in the Supplemental Information.

### Imaging based high-throughput screen

To screen for modifiers of KCNQ1 trafficking, expression of WT mycKCNQ1-mEGFP (used for compound testing) or E115G mycKCNQ1-mEGFP (used as a control) was induced by treating the respective stable cell line with 1 µg/mL doxycycline 24 hours before plating cells into 384-well microscopy plates (Greiner, 781090) coated with PLL. Compounds used for HTS were obtained from the Vanderbilt High Throughput Screening core facility dissolved in DMSO at 10 mM. Four to six hours after plating, compounds were diluted in media with doxycycline and added to wells to a final concentration of 10 µM (0.1% DMSO). 0.1% DMSO was used as a control. Cells were incubated with compounds for ∼16 hours before staining. The population of mycKCNQ1-mEGFP at the cell surface was labeled by incubating cells with an anti-myc mouse antibody (Cell Signaling, 2276S) diluted 1:500 in media without pen/strep and FBS for 30 minutes at room temperature. Cells were then fixed with 4% paraformaldehyde (Santa Cruz Biotechnology, sc-281692) for 30 minutes. After fixing, cells were washed three times, with phosphate buffered saline with calcium and magnesium (PBS++) (Sigma, D1283). Cells were then incubated with an anti-mouse AlexaFluor647 conjugated secondary antibody (Cell Signaling, 4410S) diluted 1:1000 in antibody dilution buffer (PBS++ with 5% goat serum (Gibco,16210-064)) for 45 minutes. Cells were again washed 3 times with PBS++ then permeabilized with PBS++ with 0.3% Triton-X and 0.1% BSA for 15 minutes. Cell membranes were then stained with CF®405S-conjugated wheat germ agglutinin (WGA) (Biotium, 29027) at a 1:200 dilution in PBS++ for 30 - 45 minutes. Lastly, cells were washed 3 times with PBS++ and imaged using an ImageXpress Micro Confocal high content imager (described below).

### Compounds screened

All screened small molecules were purchased from commercial sources, stored frozen in DMSO at a concentration of 10 mM, and managed by the Vanderbilt High-Throughput Screening Facility. Small molecules from the following libraries were screened:

- FDA approved drug collection (Selleck Chemicals, Houston, TX).
- Vanderbilt discovery library (purchased from Life Chemicals, Ontario, Canada). Curated list of ∼100,000 compounds selected by Vanderbilt scientists to represent maximum chemical diversity and minimal pan-assay interference. The first 20,000 compounds of this library, which represent the chemical diversity of the full ∼100,000 compounds in the library, were screened in this work.
- Bioactive lipids library (Cayman Chemical, Ann Arbor, MI).
- Steroid-like compound library (ChemDiv, San Diego, CA).

After initial screening, fresh VU0494372 (product number F1562-0024) powder was purchased from Life Chemicals and was used for all subsequent experiments.

### High content imaging

Imaging of 384-well plates containing LLP-int WT (or E115G, in control wells) KCNQ1-mEGFP cells treated with DMSO or screening compounds was performed using an Image Xpress Micro Confocal (IXMC) high content imaging system (Molecular Devices, San Jose, CA). Cell surface KCNQ1, total KCNQ1, and KCNQ1 trafficking ratio for each well were quantified using a custom module designed in the IXMC’s image analysis software - MetaXpress. Details of imaging and microscope settings, as well as a description of the MetaXpress custom module are described in the Supplemental Information.

### Hit picking and quality control

Data obtained from the custom analysis module was visualized and analyzed using the Vanderbilt High-Throughput screening core facility’s data visualization system Waveguide in conjunction with BIOVIA Pipeline Pilot (Dassault Systemes, Paris, France). A robust Z score:

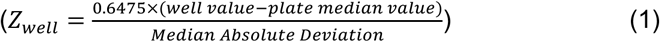

was calculated for each compound screened. Hits were defined as compounds with a robust Z score of >3 or <-3. Additional quality control metrics were included to flag cells with undesirable characteristics, like toxicity or aggregation. Wells where cell counts were greater than 5 standard deviations below the average cell count in the plate, and wells where the “laser focus score”, a metric outputted by the IXMC that quantifies how well the instrument was able to focus on a well, was lower than 20, were flagged and eliminated from hit lists. Finally, quality control metrics (Z’ and percent coefficient of variance (%CV), equations shown in Supplemental Information) were calculated for each plate, to ensure the robustness of the screen and quality of each individual plate.

During hit confirmation experiments, the same values were calculated for each compound for each biological replicate. Hits were considered “confirmed” if they met the Z score threshold of >3 or <-3 on at least 2 of the 3 biological replicates.

### Flow cytometry-based trafficking assay

Trafficking of KCNQ1 in the LLP-int WT mycKCNQ1 cell line was quantified using a previously published flow cytometry-based trafficking assay (13, 21). In short, KCNQ1 expression was induced for ∼24 hours, cells were treated with compound or DMSO control ∼16 hours, and then cell surface and internal populations of KCNQ1 were stained. Cells were washed once with PBS (Corning) plus 0.2% sodium azide (PBS-FC), detached with a solution of PBS-FC plus 0.5 mM EDTA and 0.5% bovine serum albumin (BSA) (Fisher Bioreagents, BP9704-100), and pelleted by centrifugation. Cell surface mycKCNQ1 was stained by incubating cells in PBS-FC plus 5% FBS (PBS-FC +FBS) containing anti-myc phycoerythrin (PE) (Cell Signaling, 3739S) or AlexaFluor 488-(Cell Signaling, 2279S) conjugated antibody at a 1:100 dilution for 30 minutes. Cells were then fixed by adding Fix & Perm Medium A (Invitrogen, GAS004) for 15 minutes. Cells were washed twice, then permeabilized while simultaneously labeling internal mycKCNQ1 with Fix & Perm Medium B (Invitrogen, GAS004) with anti-myc AlexaFluor 647-conjugated antibody (Cell Signaling, 2233S), diluted 1:100, for 30 minutes. Cells were again washed 2 times and then resuspended in PBS-FC + FBS for flow cytometry. The following controls were also prepared: untransfected (or uninduced) and unstained cells, single color controls (total protein stained with one antibody), and background staining control (uninduced or empty vector-transfected cells stained with antibodies) for use in flow cytometry analysis.

### Flow cytometry

Flow cytometry analysis was conducted using a 5-laser BD LSRFortessa flow cytometer (BD Biosciences, Franklin Lanes, NJ). Compensation was determined using unstained cells and single-color controls described above. Gates were drawn to select single cells that were mCherry (for LLP-int cell lines) or EGFP (for transiently transfected cells) positive. AlexaFluor647 and AlexaFluor 488 (LLP-int cell lines) or PE (transiently transfected cells) fluorescence intensities were then quantified for 2,500 cells per sample. Single cell data was exported using FlowJo data analysis software (Ashland, OR) and used for calculating average cell surface KCNQ1, total KCNQ1, and KCNQ1 trafficking efficiency. Details of background subtraction and brightness correction calculations for cell surface KCQN1 and total KCNQ1, as well as KCNQ1 trafficking efficiency calculations are described in the Supplemental Information.

### RNA extraction, cDNA generation, and qPCR

Relative KCNQ1 and GAPDH (control) mRNA levels were determined using RT-qPCR. Expression of mycKCNQ1 in the LLP-int WT mycKCNQ1 cell line was induced by treatment with 1 µg/mL doxycycline for 24 hours, then cells were treated with 0.1% DMSO or 10 µM VU0494372 for 16 hours. RNA was extracted and purified from cells using the Cold Spring Harbor “Purification of Total RNA from Mammalian Cells and Tissues” protocol (69). RNA concentration and A260/A280 were determined with a Thermoscientific NanoDrop Lite Spectrophotometer (Waltham, MA). RNA was then reverse transcribed into cDNA using the Invitrogen SuperScript IV VILO Master Mix kit (Invitrogen, 11766050). For qPCR experiments, diluted cDNA and qPCR primers were combined with FAST SYBR qPCR master mix (Applied biosystems, 4385612) and plated in triplicate in a qPCR clear reaction plate (Applied Biosystems, A36924). Details of qPCR primers, thermal cycling protocol, and calculation of fold change are described in detail in the Supplemental Information.

### Cycloheximide chase assay

WT mycKCNQ1 expression was induced in LLP-int cells by adding doxycycline to the media 24 hours before additional treatment with either 0.1% DMSO or 10 µM VU0494372 for 16 hours. To assess rate of KCNQ1 degradation, cells were then treated with either 50 µg/mL cycloheximide (CHX) (Cell Signaling, 2112S) or 0.1% DMSO (as a control) for 0, 3, 6, or 9 hours. Soluble KCNQ1 was collected using the same lysis method as CETSA experiments. Total protein in lysates was quantified using Pierce 660 nm protein assay reagent (Thermo Scientific, 22660), and 6-7 µg protein was loaded per well into SDS PAGE gels. Western blotting was performed as described in Supplemental Information for CETSA experiments, except blots were cut above the 50 kDa ladder band and the top half was probed with an anti-KCNQ1 C-terminal domain antibody (Alomone Labs, APC-022) and the bottom half was probed with an anti β-actin antibody (Cell Signaling, 8457S) as a loading control. Bands corresponding to mycKCNQ1 (approximately 70 kDa) and β-actin were quantified with the Fiji/ImageJ BandPeakQuantification plugin (70, 71), and mycKCNQ1 band intensities were normalized to the β-actin intensity in the same lane. Normalized mycKCNQ1 band intensities were then represented as a “percent remaining” from the 0 hour treatment time point and plotted using GraphPad Prism 10.

### Cellular thermal shift assay (CETSA)

The effects of compounds on the thermal stability of KCNQ1 in cells was assessed with a Cellular Thermal Shift Assay (CETSA). The principles and protocol for CETSA has been described in detail in several publications (72, 73), and CETSA has also been optimized for studies of membrane proteins more broadly (74), and for studying KCNQ1 in particular (12). Briefly, LLP-int WT mycKCNQ1-expressing cells were treated with 0.2% DMSO or 15 µM VU0494372 for 16 hours, then detached and collected. Cells were diluted with PBS to a concentration of 1 x 10^7^ cells/mL and protease inhibitor cocktail (Sigma-Aldrich, P8340) was added at a 1:100 dilution. Cells were then divided into 50 µL aliquots in PCR tubes and heated to the following temperatures: 37 °C (unheated control), 40 °C, 44 °C, 48 °C, 52 °C, 56 °C, 60 °C, 64 °C, 68 °C, 72 °C, and 76 °C for three minutes using a thermal cycler. After heat shock, 2 µL of 25% digitonin (Sigma, D141) in water was added to each PCR tube, and cells were lysed with three rounds of freeze-thaw with liquid nitrogen. The insoluble fraction was pelleted with centrifugation and supernatant was transferred to a fresh tube containing TCEP.

Soluble KCNQ1 from CETSA experiments was then quantified with western blotting. Details are described in the Supplemental Information. A rabbit IgG anti-KCNQ1 C-terminal domain antibody (Alomone Labs, APC-022) diluted 1:2500 in blocking buffer and a anti-rabbit IgG/HRP conjugated antibody (Cell Signaling, 7074S) diluted 1:5000 in blocking buffer were used to probe KCNQ1 bands. Western blots were imaged by treating with Clarity Western ECL substrate (Bio-Rad, 170-5060) and detecting chemiluminescence. Bands corresponding to mycKCNQ1 (approximately 70 kDa) were quantified with the Fiji/imageJ BandPeakQuantification plugin (70, 71). Band intensities were normalized to the 37 °C sample band within each western blot. IC50 (T_Agg_) values were calculated for each replicate and were plotted to compare the DMSO and compound-treated curves.

### Cell viability

Viability of cells after treatment with 0.1% - 0.2% DMSO or 10 - 30 µM VU0494372 for 16 hours was assessed with trypan blue staining. Cells were detached with trypsin, retaining all media to ensure collection of all cells. Cell solutions were diluted 1:2 in trypan blue (Invitrogen, T10282) and then density and viability (percent live cells) were determined with a Countess 3 FL (Invitrogen) automated cell counter. Cells killed with ∼50% ethanol and were assessed for viability as a control.

### Automated patch clamp recording

#### Plasmids and site-directed mutagenesis

Full length cDNA encoding wild type human (WT) KCNQ1 (GenBank accession number AF000571) were engineered in the mammalian expression vector pIRES2-EGFP (BD Biosciences-Clontech, Mountain View, CA, USA) as previously described (67). This vector enabled expression of untagged channel subunits with fluorescent proteins as a means for tracking successful cell transfection.

#### Electroporation

Plasmid encoding WT or mutant KCNQ1 was transiently transfected into Chinese Hamster Ovary (CHO) cells stably transfected with human KCNE1 (CHO-KCNQ1 cells) by electroporation using the Maxcyte STX system (MaxCyte Inc., Gaithersburg, MD, USA) to generate WT I_Ks_-expressing cells, as previously described (67). Electroporated cells were frozen in 1 mL aliquots at 1.8×10^6^ viable cells/mL and stored in liquid nitrogen until they were used in experiments.

#### Electrophysiology

The day before automated patch clamp recording, electroporated cells were thawed. For analysis of chronic exposure of I_Ks_ cells to VU0494372, cells were grown for 10 hrs., and then exposed to various concentrations of VU0494372 or vehicle (DMSO) for 16 hrs. For analysis of acute exposure to VU0494372, cells were grown for 26 hrs. Prior to experiments, cells were harvested using 0.25% trypsin. Aliquots were used to determine cell number and viability by automated cell counting and transfection efficiency by flow cytometry. Cells were then diluted to 300,000 cells/mL with external bath solution (see below) and allowed to recover 60 minutes at 15 °C while shaking on a rotating platform. Chronic exposure cells were continuously exposed to VU0494372 until the cell harvest step, during which compound was washed away and after which compound was no longer added. For acute exposure to VU0494372, untreated cells were exposed to various concentrations of VU0494372 or vehicle (DMSO) prior to attaining whole-cell configuration and currents recorded ∼5 minutes after initiating exposure. Automated patch clamp experiments were performed using the Syncropatch 384 platform (Nanion Technologies, Munich, Germany) and 4-hole, 384-well S-Type (0.75-1.25 MΩ in control solutions) recording chips. Pulse generation and data collection were carried out with PatchController384 V.1.3.0 and DataController384 V1.2.1 software (Nanion Technologies). Whole-cell currents were recorded at room temperature in the whole-cell configuration, filtered at 3 kHz and acquired at 10 kHz. The access resistance and apparent membrane capacitance were estimated using built-in protocols. The external bath solution contained 140 mM NaCl, 4 mM KCl, 2 mM CaCl_2_, 1 mM MgCl_2_, 10 mM HEPES, 5 mM glucose, at pH 7.4. The internal solution contained 60 mM KF, 50 mM KCl, 10 mM NaCl, 10 mM HEPES, 10 mM EGTA, 2 mM ATP-Mg, at pH 7.2. Whole-cell currents were elicited from a holding potential of -80 mV using 2000 ms depolarizing pulses (from -80 mV to +60 mV in +10mV steps, every 10 secs) followed by a 2000 ms step to -30 mV to analyze tail currents and channel deactivation rate. Non-specific currents were eliminated by recording whole-cell currents before and after addition of the *I*_Ks_ selective blocker JNJ-303 (2 μM; Tocris Bioscience, Minneapolis, MN). Only JNJ-303 sensitive currents and recordings with seal resistance ≥ 0.1 GΩ and series resistance < 10 MΩ were analyzed.

Data were analyzed and plotted as previously described (67). Whole-cell currents were normalized for membrane capacitance. Peak currents were recorded at 1990 ms after the start of the voltage pulse. The voltage-dependence of activation was calculated by fitting the normalized G-V curves with a Boltzmann function (tail currents measured at -30 mV).

### Computational binding site prediction of VU0494372

Computational docking was used to identify potential binding sites of VU0494372 to KCNQ1. An initial conformer of VU0494372 was obtained in Structured Data Files (SDF) format from the Vanderbilt Discovery Collection compound library managed by the Vanderbilt High Throughput Screening Core. VU0494372 was first blind docked to two different KCNQ1 conformations from published structural models: PDB 8SIK (48) and PDB 7XNK (49). Only the KCNQ1 tetramer was used (without calmodulin or any ions), including transmembrane and cytosolic domains. We separately blind docked VU0494372 to each KCNQ1 construct using DiffDock, (50). Seven of the ten poses predicted by DiffDock using 8SIK placed VU0494372 at a site partially overlapping with the known docking site of ML277. All of the predicted poses generated by DiffDock using 7XNK place VU0494372 inside the pore domain, likely due to the large pocket formed in the open state of the pore. Using the docked poses generated by DiffDock with PDB 8SIK as starting positions, we applied a thorough targeted docking procedure to refine and score the poses. We used the BioChemicalLibrary ConformerGenerator (75) function to perform 2000 conformer generation iterations and select the top 100 conformers. We generated a Rosetta parameters file and associated PDB files of the compound conformers and used the RosettaLigand (51, 52) small molecule docking protocol to generate 200 bound structural models for each of the DiffDock starting positions. We used the Score12 score function with the restore_pre_talaris_2013_behavior flag as recommended for small molecule scoring (76). RosettaLigand calculates several protein- and ligand-specific energy score terms and combines them to determine an overall docking score (51). A more negative ligand interface score indicates a more energetically favorable binding energy. The most energetically favorable docked poses predict interactions with residues adjacent to those in the experimentally determined ML277 binding pocket.

### Re-synthesis of VU0608712 and synthesis of analogs

Synthesis and characterization of 1-(4-methylpiperazin-1-yl)-3-(4-(2,4,4-trimethylpentan-2-yl)phenoxy)propan-2-ol (VU0494372), 1-morpholino-3-(4-(2,4,4-trimethylpentan-2-yl)phenoxy)propan-2-ol (VU0489336), and 1-(piperidin-1-yl)-3-(4-(2,4,4-trimethylpentan-2-yl)phenoxy)propan-2-ol (VU0963991) are described in the Supplemental Information. Solvents were obtained from commercial sources. Commercial reagents were used as received. The compounds were purified by Teledyne ISCO normal phase column chromatograph system. Where necessary, preparative reverse phase HPLC was conducted on a Gilson HPLC system using a Phenomenex Luna column (100 Å, 50 x 21.20 mm, 5 μm C18) with UV/Vis detection. ^1^H NMR spectra were recorded on Bruker 400 MHz spectrometer and are reported relative to deuterated solvent signals. Data for ^1^H NMR spectra are reported as follows: chemical shift (δ ppm), multiplicity (s = singlet, d = doublet, t = triplet, q = quartet, quint = quintet, m = multiplet, br = broad, app = apparent), coupling constants (Hz), and integration. ^13^C NMR spectra were recorded on Bruker 100, 125, or 150 MHz spectrometers and are reported relative to deuterated solvent signals. LC/MS was conducted and recorded on an Agilent Technologies 6130 Quadrupole instrument. HRMS was conducted and recorded at the Notre Dame Mass Spectrometry and Proteomics Facility on a micrOTOF II instrument.

### Statistics

All statistical tests performed on data included in this publication were performed using GraphPad Prism version 8 or version 10. Normally distributed data was analyzed with an unpaired t-test when 2 values were compared, and a one-way ANOVA with Dunnett’s multiple comparisons test for follow up when 3 or more values were compared. When data was not normally distributed (in the case of data normalized to the control value, for example), a Wilcoxon matched-pairs signed rank test was performed 2 values were compared, and a Kruskal-Wallis test with Dunn’s multiple comparison test for follow up was performed when 3 or more values were compared. A P-value less than 0.05 was considered significant. All graphs show data mean and error bars represent standard deviation.

### Data availability

All data values for all graphs included in this paper are reported in the Supporting Data File. All data and materials are available upon request. The custom multiple sequence alignment tool Multiple Sequence Iterative Comparator (MuSIC) used for analysis of some DNA sequencing results is available at https://doi.org/10.18131/h6hc6-n0j20.

## Author Contributions

Project conceptualization: KRCM, ALG, CRS; Experiment/methods design: KRCM, CGV, ACCG, IMR, KMS, JAB, TPH, KRB, GAS, AGW, KVL, ALG, CRS; Performed experiments: KRCM, CGV, ACCG, IMR, MCW, TPH, RRD; Data analysis/interpretation: KRCM, CGV, ACCG, IMR, KMS, MCW, JAB, TPH, ELD, GAS, AGW, KVL, ALG, CRS; Supervision: JM, ALG, and CRS; Writing: KRCM and CRS; Editing: KRCM, CGV, ACCG, IMR, KMS, MCW, TPH, KRB, AGW, JM, KVL, ALG, and CRS.

## Funding Support

Funding for this work comes from the following sources: National Institutes of Health grant 4T32GM065086-14 (KRCM), National Institutes of Health grant F31HL168964-01 (KRCM), National Institutes of Health grant R01 HL122010 (ALG and CRS), National Institutes of Health grant T32 DK007061 (ACCG), Deutsche Forschungsgemeinschaft (DFG, German Research Foundation) SFB1423, project number 421152132 (JM), Humboldt Professorship of the Alexander von Humboldt Foundation (JM), Vanderbilt Institute of Chemical Biology (IMR, JAB, TPH, ELD, GAS, AGW, HTS core facility, VMC Flow Cytometry Shared Resource), Vanderbilt Ingram Cancer Center P30 CA68485 (HTS Core Facility), National Institutes of Health grant NCI R50CA211206 (JAB), Vanderbilt Digestive Disease Research Center DK058404 (VMC Flow Cytometry Shared Resource)

## Supporting information

Supplemental Methods, Figures, and References

## Acknowledgments

We would like to thank Kenneth Matreyek for sharing the “LLP-int” cells and AttB shuttle vector used to generate stable cell lines used in these studies and Arina Hadziselimovic for her assistance with product ordering, primer design, and management of day-to-day lab activities. Flow Cytometry experiments were performed in the VMC Flow Cytometry Shared Resource. Experiments were performed in the Vanderbilt High-Throughput Screening (V-HTS) Core Facility with compound management assistance provided by Corbin Whitwell. Imaging of Western blots reported in this publication was supported by the Vanderbilt University Cellular and Developmental Biology Resource Core. Chemical synthesis and compound analysis was provided by the Vanderbilt Institute of Chemical Biology Molecular Design and Synthesis Center. Panels A and B in Figure 1 were made using BioRender (https://www.biorender.com). Marvin was used for drawing and displaying chemical structures, Marvin 21.1.79, Chemaxon (https://www.chemaxon.com).

## Notes

### Competing Interest Statement

Alfred George serves on the Scientific Advisory Board of Tevard Biosciences. Alfred George received grant funding from Biohaven Pharmaceuticals.

## References

1. Schwartz PJ, Periti M, Malliani A. The long Q-T syndrome. American Heart Journal. 1975;89(3):378– 390.

2. Crotti L, et al. Congenital long QT syndrome. Orphanet J Rare Dis. 2008;3(1):18.

3. Schwartz PJ, et al. Prevalence of the congenital long-QT syndrome. Circulation. 2009;120(18):1761– 1767.

4. Apgar TL, Sanders CR. Compendium of causative genes and their encoded proteins for common monogenic disorders. Protein Science. 2022;31(1):75–91.

5. Schwartz PJ, MD LC, Insolia R. Long QT Syndrome: From Genetics to Management. Circ Arrhythm Electrophysiol. 2012;5(4):868–877.

6. Wang Q, et al. Positional cloning of a novel potassium channel gene: KVLQT1 mutations cause cardiac arrhythmias. Nat Genet. 1996;12(1):17–23.

7. Abbott GW. Biology of the KCNQ1 Potassium Channel [Internet]. New Journal of Science. 2014. 10.1155/2014/237431.

8. Brewer KR, et al. Structures Illuminate Cardiac Ion Channel Functions in Health and in Long QT Syndrome. Front Pharmacol. 2020;11. 10.3389/fphar.2020.00550.

9. Moss AJ, et al. Clinical aspects of type-1 long-QT syndrome by location, coding type, and biophysical function of mutations involving the KCNQ1 gene. Circulation. 2007;115(19):2481–2489.

10. Vincent GM, et al. High Efficacy of β-Blockers in Long-QT Syndrome Type 1: Contribution of Noncompliance and QT-Prolonging Drugs to the Occurrence of β-Blocker Treatment “Failures.” Circulation. 2009;119(2):215–221.

11. Chockalingam P, et al. Not All Beta-Blockers Are Equal in the Management of Long QT Syndrome Types 1 and 2. Journal of the American College of Cardiology. 2012;60(20):2092–2099.

12. Brewer KR, et al. Integrative analysis of KCNQ1 variants reveals molecular mechanisms of type 1 long QT syndrome pathogenesis. Proc Natl Acad Sci USA. 2025;122(8):e2412971122.

13. Huang H, et al. Mechanisms of KCNQ1 channel dysfunction in long QT syndrome involving voltage sensor domain mutations. Science Advances. 2018;4(3):eaar2631.

14. McKeone R, et al. Assessing the correlation between mutant rhodopsin stability and the severity of retinitis pigmentosa. Mol Vis. 2014;20:183–199.

15. Athanasiou D, et al. The molecular and cellular basis of rhodopsin retinitis pigmentosa reveals potential strategies for therapy. Progress in Retinal and Eye Research. 2018;62:1–23.

16. Hollingsworth TJ, Gross AK. Defective Trafficking of Rhodopsin and Its Role in Retinal Degenerations. International Review of Cell and Molecular Biology. Elsevier; 2012:1–44.

17. Kaushal S, Khorana HG. Structure and Function in Rhodopsin. 7. Point Mutations Associated with Autosomal Dominant Retinitis Pigmentosa. Biochemistry. 1994;33(20):6121–6128.

18. Sung CH, et al. Functional heterogeneity of mutant rhodopsins responsible for autosomal dominant retinitis pigmentosa. Proc Natl Acad Sci USA. 1991;88(19):8840–8844.

19. Mendre C, Mouillac B. Misfolding of vasopressin receptors: biased agonist pharmacochaperones as potential therapeutics. Advances in Protein Chemistry and Structural Biology. Elsevier; 2019:249–272.

20. Ulloa-Aguirre A, Janovick JA. Modulation of proteostasis and protein trafficking: a therapeutic avenue for misfolded G protein-coupled receptors causing disease in humans. Emerging Topics in Life Sciences. 2019;3(1):39–52.

21. Schlebach JP, et al. Conformational Stability and Pathogenic Misfolding of the Integral Membrane Protein PMP22. J Am Chem Soc. 2015;137(27):8758–8768.

22. Roland Naef, Suter U. Impaired Intracellular Trafficking Is a Common Disease Mechanism ofPMP22Point Mutations in Peripheral Neuropathies. Neurobiology of Disease. 1999;6(1):1–14.

23. Li J, et al. The PMP22 Gene and Its Related Diseases. Mol Neurobiol. 2013;47(2):673–698.

24. Cheung JC, Deber CM. Misfolding of the cystic fibrosis transmembrane conductance regulator and disease. Biochemistry. 2008;47(6):1465–1473.

25. Kerem B, et al. Identification of the cystic fibrosis gene: genetic analysis. Science. 1989;245(4922):1073–1080.

26. Morales MM, Capella MA, Lopes AG. Structure and function of the cystic fibrosis transmembrane conductance regulator. Braz J Med Biol Res. 1999;32(8):1021–1028.

27. Jilling T, Kirk KL. The Biogenesis, Traffic, and Function of the Cystic Fibrosis Transmembrane Conductance Regulator. International Review of Cytology. Elsevier; 1997:193–241.

28. Southern KW, et al. Correctors (specific therapies for class II CFTR mutations) for cystic fibrosis. Cochrane Database Syst Rev. 2018;8:CD010966.

29. Ohgane K, Dodo K, Hashimoto Y. Retinobenzaldehydes as proper-trafficking inducers of folding-defective P23H rhodopsin mutant responsible for Retinitis Pigmentosa. Bioorganic & Medicinal Chemistry. 2010;18(19):7022–7028.

30. Noorwez SM, et al. Retinoids Assist the Cellular Folding of the Autosomal Dominant Retinitis Pigmentosa Opsin Mutant P23H. Journal of Biological Chemistry. 2004;279(16):16278–16284.

31. Chen Y, et al. A novel small molecule chaperone of rod opsin and its potential therapy for retinal degeneration. Nat Commun. 2018;9(1):1976.

32. Van Goor F, et al. Correction of the F508del-CFTR protein processing defect in vitro by the investigational drug VX-809. Proc Natl Acad Sci USA. 2011;108(46):18843–18848.

33. Mouillac B, Mendre C. Vasopressin receptors and pharmacological chaperones: From functional rescue to promising therapeutic strategies. Pharmacological Research. 2014;83:74–78.

34. Mouillac B, Mendre C. Pharmacological Chaperones as Potential Therapeutic Strategies for Misfolded Mutant Vasopressin Receptors. In: Ulloa-Aguirre A, Tao Y-X, eds. Targeting Trafficking in Drug Development. Cham: Springer International Publishing; 2017:63–83.

35. Castellani C, et al. Consensus on the use and interpretation of cystic fibrosis mutation analysis in clinical practice. Journal of Cystic Fibrosis. 2008;7(3):179–196.

36. Middleton PG, et al. Elexacaftor–Tezacaftor–Ivacaftor for Cystic Fibrosis with a Single Phe508del Allele. N Engl J Med. 2019;381(19):1809–1819.

37. Baroni D. Unraveling the Mechanism of Action, Binding Sites, and Therapeutic Advances of CFTR Modulators: A Narrative Review. CIMB. 2025;47(2):119.

38. Van Goor F, et al. Rescue of CF airway epithelial cell function in vitro by a CFTR potentiator, VX-770. Proc Natl Acad Sci USA. 2009;106(44):18825–18830.

39. Highlights of prescribing information, Trikafta. https://www.accessdata.fda.gov/drugsatfda_docs/label/2023/217660s000lbl.pdf. Accessed September 16, 2025.

40. Bihler H, et al. In vitro modulator responsiveness of 655 CFTR variants found in people with cystic fibrosis. Journal of Cystic Fibrosis. 2024;23(4):664–675.

41. Mattmann ME, et al. Identification of (R)-N-(4-(4-methoxyphenyl)thiazol-2-yl)-1-tosylpiperidine-2-carboxamide, ML277, as a novel, potent and selective Kv7.1 (KCNQ1) potassium channel activator. Bioorg Med Chem Lett. 2012;22(18):5936–5941.

42. Bohannon BM, et al. Mechanistic insights into robust cardiac IKs potassium channel activation by aromatic polyunsaturated fatty acid analogues. eLife. 2023;12:e85773.

43. Salata JJ, et al. A Novel Benzodiazepine that Activates Cardiac Slow Delayed Rectifier K+ Currents. Molecular Pharmacology. 1998;54(1):220–230.

44. Zheng Y, et al. Hexachlorophene Is a Potent KCNQ1/KCNE1 Potassium Channel Activator Which Rescues LQTs Mutants. PLoS ONE. 2012;7(12):e51820.

45. Matreyek KA, et al. An improved platform for functional assessment of large protein libraries in mammalian cells. Nucleic Acids Res. 2020;48(1):e1.

46. Kanki H, et al. A Structural Requirement for Processing the Cardiac K+ Channel KCNQ1. Journal of Biological Chemistry. 2004;279(32):33976–33983.

47. Wilson AJ, et al. Abnormal KCNQ1 trafficking influences disease pathogenesis in hereditary long QT syndromes (LQT1). Cardiovascular Research. 2005;67(3):476–486.

48. Mandala VS, MacKinnon R. The membrane electric field regulates the PIP _2_ -binding site to gate the KCNQ1 channel. Proc Natl Acad Sci USA. 2023;120(21):e2301985120.

49. Ma D, et al. Structural mechanisms for the activation of human cardiac KCNQ1 channel by electro-mechanical coupling enhancers. Proc Natl Acad Sci USA. 2022;119(45):e2207067119.

50. Corso G, et al. DiffDock: Diffusion Steps, Twists, and Turns for Molecular Docking [preprint]. 2022. 10.48550/ARXIV.2210.01776.

51. Lemmon G, Meiler J. Rosetta Ligand Docking with Flexible XML Protocols. In: Baron R, ed. Computational Drug Discovery and Design. New York, NY: Springer New York; 2012:143–155.

52. Meiler J, Baker D. ROSETTALIGAND: Protein–small molecule docking with full side-chain flexibility. Proteins. 2006;65(3):538–548.

53. Boulmpou A, et al. The Uncommon Phenomenon of Short QT Syndrome: A Scoping Review of the Literature. JPM. 2025;15(3):105.

54. Chen Y-H, et al. KCNQ1 gain-of-function mutation in familial atrial fibrillation. Science. 2003;299(5604):251–254.

55. Seebohm G, et al. Molecular Determinants of KCNQ1 Channel Block by a Benzodiazepine. Molecular Pharmacology. 2003;64(1):70–77.

56. Bleich M, et al. K V LQT channels are inhibited by the K + channel blocker 293B. Pfl□gers Archiv European Journal of Physiology. 1997;434(4):499–501.

57. Dilly KW, et al. Overexpression of β2-Adrenergic Receptors cAMP-dependent Protein Kinase Phosphorylates and Modulates Slow Delayed Rectifier Potassium Channels Expressed in Murine Heart. Journal of Biological Chemistry. 2004;279(39):40778–40787.

58. Thompson E, et al. cAMP-dependent regulation of *IKs* single-channel kinetics. Journal of General Physiology. 2017;149(8):781–798.

59. Cui C, et al. Mechanisms of KCNQ1 gating modulation by KCNE1/3 for cell-specific function [preprint]. 2025. 10.1101/2025.06.08.658478.

60. Wickenden AD, et al. Retigabine, A Novel Anti-Convulsant, Enhances Activation of KCNQ2/Q3 Potassium Channels. Mol Pharmacol. 2000;58(3):591–600.

61. Song Y, et al. Genetic features and pharmacological rescue of novel Kv7.2 variants in patients with epilepsy. J Med Genet. 2025;62(4):231–241.

62. Hou P, et al. ML277 specifically enhances the fully activated open state of KCNQ1 by modulating VSD-pore coupling. eLife. 2019;8:e48576.

63. Veit G, et al. Allosteric folding correction of F508del and rare CFTR mutants by elexacaftor-tezacaftor-ivacaftor (Trikafta) combination. JCI Insight. 2020;5(18):e139983.

64. Xu Y, et al. Probing Binding Sites and Mechanisms of Action of an I Ks Activator by Computations and Experiments. Biophysical Journal. 2015;108(1):62–75.

65. Leidenheimer NJ, Ryder KG. Pharmacological chaperoning: A primer on mechanism and pharmacology. Pharmacological Research. 2014;83:10–19.

66. De Git KCG, et al. Cardiac ion channel trafficking defects and drugs. Pharmacology & Therapeutics. 2013;139(1):24–31.

67. Vanoye CG, et al. High-Throughput Functional Evaluation of KCNQ1 Decrypts Variants of Unknown Significance. Circ Genom Precis Med. 2018;11(11):e002345.

68. Matreyek KA, Stephany JJ, Fowler DM. A platform for functional assessment of large variant libraries in mammalian cells. Nucleic Acids Research. 2017;45(11):e102–e102.

69. Green MR, Sambrook J. Purification of Total RNA from Mammalian Cells and Tissues. Cold Spring Harb Protoc. 2020;2020(1):pdb.prot101659.

70. Schindelin J, et al. Fiji: an open-source platform for biological-image analysis. Nat Methods. 2012;9(7):676–682.

71. Ohgane K, Yoshioka H. Quantification of Gel Bands by an Image J Macro, Band/Peak Quantification Tool v1 [preprint]. 2019. 10.17504/protocols.io.7vghn3w.

72. Jafari R, et al. The cellular thermal shift assay for evaluating drug target interactions in cells. Nat Protoc. 2014;9(9):2100–2122.

73. Martinez Molina D, Nordlund P. The Cellular Thermal Shift Assay: A Novel Biophysical Assay for In Situ Drug Target Engagement and Mechanistic Biomarker Studies. Annu Rev Pharmacol Toxicol. 2016;56(1):141–161.

74. Kawatkar A, et al. CETSA beyond Soluble Targets: a Broad Application to Multipass Transmembrane Proteins. ACS Chem Biol. 2019;14(9):1913–1920.

75. Brown BP, et al. Introduction to the BioChemical Library (BCL): An Application-Based Open-Source Toolkit for Integrated Cheminformatics and Machine Learning in Computer-Aided Drug Discovery. Front Pharmacol. 2022;13:833099.

76. Smith ST, Meiler J. Assessing multiple score functions in Rosetta for drug discovery. PLoS ONE. 2020;15(10):e0240450.

