## Supplemental Methods, Figures, and References for "High Throughput Screening Identifies a Small Molecule Trafficking Corrector for Long-QT Syndrome Associated KCNQ1 Variants"

<u>Supplemental Information</u>

### Materials and Methods:

#### Cell culture conditions:

All mammalian cells used in this work were cultured at 37 °C and 5% $CO_2$. HEK 293T “LLP-int” cells, and all stable cell lines generated from LLP-int cells, were cultured in high glucose Dulbecco’s Modified Eagle Medium without sodium pyruvate (Gibco, 11965-092) supplemented with 100 U/mL penicillin/streptomycin (Gibco, 15140-122) and 10% tetracycline negative FBS (Corning, 35-075-CV). LLP-int cells were grown in flasks coated with poly-L-lysine (Millipore, A-005-C). HEK 293 cells were cultured in high glucose Dulbecco’s Modified Eagle Medium without sodium pyruvate (Gibco, 11965-092) containing 100 U/mL penicillin/streptomycin (Gibco, 15140-122) and 10% FBS (Gibco, 26140-079). CHO-K1 cells constitutively expressing human KCNE1 (designated CHO-KCNE1 cells) used for electrophysiology experiments were generated as previously described (1) using the FLP-in$^{TM}$ system (Thermo Fisher Scientific, Waltham, MA, USA) and were grown in F-12 Ham nutrient medium (Gibco/Invitrogen, San Diego, CA, USA) supplemented with 10% fetal bovine serum (ATLANTA Biologicals, Norcross, GA, USA), penicillin (50 units/mL), streptomycin (50 µg/mL) and maintained under selection with hygromycin B (600 µg/mL).

#### KCNQ1 constructs and mutagenesis:

All experiments were conducted with “mycKCNQ1”: full-length human KCNQ1 (GenBank accession number AF000571) with a myc epitope (EQKLISEEDL) inserted after residue E146, which is in the S1-S2 extracellular loop of the KCNQ1 voltage sensing domain (2), or “mycKCNQ1-mEGFP”: mycKCNQ1 with a monomeric enhanced green fluorescent protein (mEGFP, containing the A206K mutation) fused to the C-terminus. Fusion protein generation is described below.

#### Landing pad cells:

Stable cell lines used for high-throughput screening (HTS) and subsequent experiments were generated using “LLP-int” HEK 293T cells, which were a gift from the lab of Dr. Kenneth Matreyek. Development and validation of this cell line is described in detail in previous publications (3, 4). In short, LLP-int cells contain a genomic “landing pad” DNA integration site with a tetracycline-inducible promoter upstream of an AttP integration sequence in their genome. Downstream of the AttP sequence is a blue fluorescent protein gene (used as a marker) and a Bxb1 integrase gene (which facilitates target DNA integration). Target DNA flanked by a complementary AttB site can be integrated at the AttP site when Bxb1 expression is induced by treatment with tetracycline/doxycycline. The Bxb1 and blue fluorescent protein will be pushed out of the reading frame and no longer expressed when target DNA is integrated.

**Generation of stable cell lines:**

A promotorless AttB “shuttle vector” construct (pKanAttb-KCNQ1-mEGFP-myc-IRES-mCherry-H2A-P2A-PuroR-WPRE) containing the cDNA for human WT mycKCNQ1-mEGFP, the cDNA for mCherry fused to histone H2A, and cDNA for a puromycin resistance gene, flanked by an AttB integration site, was used for generation of the main stable cell line used for high-throughput screening (SI Appendix, Fig. S1A).

First, an AttB shuttle vector containing WT mycKCNQ1 (pKan-AttB-KCNQ1-myc-IRES-mCherry-H2A-P2A-PuroR-WPRE) was generated. The open reading frame of human c-Myc tagged KCNQ1 (2) was amplified by PCR using primers 1 and 2 (Supplemental Table S1). Plasmid AttB_ACE2_IRES-mCherry-H2A-P2A-PuroR (a gift from Kenneth Matreyek; Addgene plasmid # 171594) (5) was digested with PshAI and XmaI to remove the ACE2 coding sequence then agarose gel purified using NucleoSpin Gel and PCR Clean-up (Takara). The KCNQ1 amplicon was subcloned into the PshAI / XmaI digested plasmid using NEBuilder HiFi DNA Assembly (New England Biolabs) and transformed into chemically competent Top10 cells. Selected clones were subjected to full length nanopore sequencing (Primodium labs). To enhance transgene expression and allow bacterial selection in kanamycin, the functional core of this construct encompassing the start of the AttB element through the end of the PuroR sequence was PCR amplified using primers 3 and 4 (Supplemental Table S1) then subcloned using NEBuilder HiFi DNA Assembly (New England Biolabs) into a NotI and SpeI digested plasmid (pKan-WPRE) containing a kanamycin resistance gene, a high-copy replication origin and the woodchuck hepatitis virus (WHV) posttranscriptional regulatory element (WPRE) followed by the SV40 late polyadenylation signal. All PCR reactions to this point were performed using the Q5 Hot Start High-Fidelity 2X Master Mix (New England Biolabs).

The pKan-AttB-KCNQ1-myc-IRES-mCherry-H2A-P2A-PuroR-WPRE plasmid was then used to generate a second plasmid containing a WT mycKCNQ1-mEGFP fusion protein. The mycKCNQ1 shuttle vector was amplified with primers 5 and 6 (Supplemental Table S1), and an amplicon containing the sequence for mEGFP (EGFP – GenBank Accession number PV533970.1 - with a A206K mutation to prevent dimerization) was amplified from a pEGFP-C99-N1 (Addgene) plasmid with primers 7 and 8 (Supplemental Table S1). The vector and mEGFP insert were subcloned using NEBuilder HiFi DNA Assembly (New England Biolabs). The assembly product (pKanAttb-KCNQ1-mEGFP-myc-IRES-mCherry-H2A-P2A-PuroR-WPRE) was transformed into XL1 Blue chemically competent e. coli, purified with a QIAprep Spin (Qiagen) miniprep kit, and sequenced with Sanger Sequencing (Genewiz/Azenta) to confirm generation of the correct fusion product.

An additional shuttle vector with E115G in mycKCNQ1-mEGFP was generated using QuikChange mutagenesis (Agilent) and primers 9 and 10 (Supplemental Table S1). Variations V207M, G189E, and G179S were introduced to the mycKCNQ1 (non-fusion) shuttle vector with primers 11-16 (Supplemental Table S1). Sanger sequencing (Genewiz/Azenta) was used to confirm generation of desired mutation.

LLP-int stable cell lines expressing mycKCNQ1-mEGFP were generated by adding 1 μg/mL doxycycline (Acros Organics, 446060050) to the LLP-int cell culture media for 24 hours to induce Bxb1 recombinase expression, then transfecting with the WT mycKCNQ1-mEGFP AttB shuttle vector using Fugene 6 (Promega, F6-1000) (4 μL Fugene: 2 μg DNA) diluted in Opti-MEM + GlutaMAX (Gibco, 51985-034). 24 hours after transfection, successfully integrated cells were selected by adding 5 μg/mL puromycin (Sigma-Aldrich, P8833) to the doxycycline-containing media. After approximately 4 days of selection, during which dead cells were washed away and cells were split as needed, individual cells were sorted into single wells in 96-well plates using a 5-laser BD FACS Aria III cell sorter. These cells were expanded to generate clonal stable cell lines. Clonal line "G6" was selected for use for further experiments after confirmation of doxycycline-inducible expression of mycKCNQ1-mEGFP via western blot (SI Appendix, Fig S1B). These cells were expanded and frozen in aliquots, and aliquots were thawed and used for approximately one month of screening/experiments. Clonal stable cell lines expressing

E115G mycKCNQ1-mEGFP and WT mycKCNQ1 (no mEGFP fusion) were generated using the same method. The E115G mycKCNQ1-mEGFP cell line was used as a control in the HTS experiments, and the WT mycKCNQ1 cell line was used for flow cytometry trafficking assays, CETSA, qPCR, and other follow-up experiments.

LLP-int WT KCNQ1, G179S KCNQ1, G189E KCNQ1, and V207M KCNQ1 cell lines used in Fig. 3 were generated in a similar fashion, but cells were selected for 7 days and the single-cell sorting step was omitted. These are therefore described as "population" cell lines instead of clonal lines. Before flow cytometry-based trafficking assay experiments, cells were cultured with no puromycin or doxycycline for at least 1 week to clear any previously expressed KCNQ1.

It should be noted that KCNE1 was not co-expressed in any of the above-described LLP-int stable cell lines.

**Imaging based high-throughput screen:**

To screen for modifiers of KCNQ1 trafficking, WT mycKCNQ1-mEGFP (used for compound testing) or E115G mycKCNQ1-mEGFP (used as a control) expression was induced by adding 1 μg/mL doxycycline to the culture media to the respective stable cell line 24 hours before plating 5,400 cells/well into black-walled 384-well microscopy plates (Greiner, 781090) coated with poly-L-lysine (Millipore, A-005-C). Compounds used for HTS were obtained from the Vanderbilt High Throughput Screening core facility dissolved in DMSO at a concentration of 10 mM. Four to six hours after plating, compounds were diluted in media containing doxycycline and added into individual wells in each plate to give a final concentration of 10 μM compound (0.1% DMSO) or only 0.1% DMSO as a control. Cells were incubated with compounds for ~16 hours before staining. The population of mycKCNQ1-mEGFP at the cell surface was labeled by incubating cells with an anti-myc mouse antibody (Cell Signaling, 2276S) diluted 1:500 in media without pen/strep and FBS for 30 minutes at room temperature. Cells were then fixed with 4% paraformaldehyde (Santa Cruz Biotechnology, sc-281692) for 30 minutes. After fixing, cells were washed three times, five minutes each wash, with 1x phosphate buffered saline with calcium and magnesium (PBS++) (Sigma, D1283). Cells were then incubated with an anti-mouse AlexaFluor647 conjugated secondary antibody (Cell Signaling, 4410S) diluted 1:1000 in antibody dilution buffer (PBS++ with 5%

goat serum (Gibco,16210-064)) for 45 minutes. Cells were again washed 3 times with PBS++ then permeabilized with permeabilization buffer (PBS++ with 0.3% Triton-X and 0.1% BSA) for 15 minutes. Cell membranes were then stained with CF®405S-conjugated wheat germ agglutinin (WGA) (Biotium, 29027) at a 1:200 dilution in PBS++ for 30 - 45 minutes. Lastly, cells were washed 3 times with PBS++ and imaged using an ImageXpress Micro Confocal high content imager (described below). All liquid handling during compound treatment and staining was performed using an Integra Biosciences Mini 96 portable electronic pipette.

**Compounds screened:**

All small molecules screened in this work were purchased from commercial sources and managed by the Vanderbilt High-Throughput Screening Facility. All compound libraries are stored frozen in DMSO at a concentration of 10 mM before distribution with a Labcyte Echo 650 or 550 Omics (Sunnyvale, CA) acoustic liquid handler. Small molecules from the following libraries were screened:

- FDA approved drug collection: ~1000 FDA-approved compounds used to treat a variety of human diseases, purchased from Selleck Chemicals (Houston, TX).
- Vanderbilt discovery library: Selected compounds from Life Chemicals (Ontario, Canada). Curated list of ~100,000 compounds were selected by Vanderbilt scientists to represent maximum chemical diversity and minimal pan-assay interference. The first 20,000 compounds of this library, which represent the chemical diversity of the full ~100,000 compounds in the library, were screened in this work.
- Bioactive lipids library: Collection of ~1000 bioactive lipids, purchased from Cayman Chemical (Ann Arbor, MI).
- Steroid-like compound library – Collection of ~500 steroid-like compound, purchased from ChemDiv (San Diego, CA).

After initial screening, fresh VU0494372 (product number F1562-0024) powder was purchased from Life Chemicals (Ontario, Canada) and was used for all subsequent experiments.

**High content imaging:**

Imaging of 384-well plates containing LLP-int WT (or E115G, in control wells) KCNQ1-mEGFP cells treated with DMSO or screening compounds was performed using an Image Xpress Micro Confocal (IXMC) high content imaging system (Molecular Devices, San Jose, CA). Images were acquired using the spinning disc confocal (60 µm pinhole) setting and the 10x Plan Apo Lambda objective (Nikon Instruments, 6500-0120). Four images were acquired per well, covering nearly the entire bottom of the 384-well plate well. The following filter sets were used: DAPI (FF01-452/45), FITC (FF01-536/40), TexasRed (FF01-624/40), and Cy5 (FF01-692/40). All lasers were set at an illumination power of 100%, and Z-offset from W1 channel was set prior to imaging each plate. Typical exposure times were as follows: DAPI ~400 ms, FITC ~500 ms, TexasRed ~200 ms, Cy5 ~750 ms. Some follow up imaging was performed with the 20X and 40X Plan Apo Lambda objectives (Nikon Instruments,1-6300-0196 and 1-6300-0412) with exposure time and other settings adjusted as needed.

Cell surface KCNQ1, total KCNQ1, and KCNQ1 trafficking ratio for each well were quantified using a custom module designed in the IXMC's image analysis software - MetaXpress (Molecular Devices, San Jose, CA). The custom module used mCherry-H2A (excited/detected with TexasRed filter set) to identify nuclei and CF®405S WGA (DAPI filter set) to identify individual cell boundaries and build a mask for each cell, and then quantified the fluorescence intensity of AlexaFluor 647 (cell surface KCNQ1, Cy5 filter set) and mEGFP (total KCNQ1, FITC filter set) within each defined mask, giving a cell-by-cell quantification of surface and total KCNQ1 in each image. The custom module then calculated a "trafficking ratio" for each cell by dividing this cell surface fluorescence value by the total fluorescence value. This "trafficking ratio" differs from the "trafficking efficiency" value calculated in later flow cytometry experiment as it does not include background fluorescence subtraction or correction for the differing brightness of the two fluorophores. The custom module also included flagging steps that prevented quantification of very small or very bright debris, which could skew quantifications. The module outputted an average value (average of each quantified cell within each well well) for each desired metric – cell surface KCNQ1, total KCNQ1, and KCNQ1 trafficking ratio. Other metrics, including average cell size (area) and number of cells quantified (cell count) were also determined for each well.

**Hit picking and quality control:**

Data obtained from the custom analysis module was visualized and analyzed using the Vanderbilt High-Throughput screening core facility's data visualization system Waveguide in conjunction with BIOVIA Pipeline Pilot (Dassault Systemes, Paris, France). A robust Z score:

$$(Z_{well} = \frac{0.6475 \times (well\ value - plate\ median\ value)}{Median\ Absolute\ Deviation}) \qquad (1)$$

was calculated for each compound screened. Hits were defined as compounds with a robust Z score of >3 or <-3. Additional quality control metrics were included to flag cells with undesirable characteristics, like toxicity or aggregation. Wells where cell counts were greater than 5 standard deviations below the average cell count in the plate, and wells where the "laser focus score", a metric outputted by the IXMC that quantifies how well the instrument was able to focus on a well, was lower than 20, were flagged and eliminated from hit lists. Finally, quality control metrics for each plate, to ensure the robustness of the screen and quality of each individual plate, were calculated. Z', a standard measure of assay quality, was calculated for each plate in the screen with the following equation:

$$Z' = 1 - \frac{3 \times (Std.Dev._{Positive\ control}) + 3 \times (Std.Dev._{Negative\ control})}{|mean_{pos.control} - mean_{neg\ control}|} \qquad (2)$$

A Z' value over 0.5 was considered good, over 0 was considered acceptable, and below zero was considered to be poor quality. A percent coefficient of variance (%CV) was also calculated for the controls in each well with the following equation:

$$\%CV = \frac{Std.Dev.}{Mean} \times 100\% \qquad (3)$$

A %CV below 10% was considered ideal and indicative of low variability within the controls, and thus a robust assay.

During hit confirmation experiments, the same values were calculated for each compound for each biological replicate. Hits were considered "confirmed" if they met the Z score threshold of >3 or <-3 on at least 2 of the 3 biological replicates.

**Flow cytometry-based trafficking assay:**

Trafficking of KCNQ1 in the LLP-int WT mycKCNQ1 cell line was quantified using a previously published flow cytometry-based trafficking assay (6, 7). KCNQ1 expression was induced for ~24 hours, and cells were treated with compound or DMSO control ~16 hours before beginning staining. Cells were washed once with phosphate buffered saline (Corning, 21-031-CV) plus 0.2% sodium azide (PBS-FC), detached with a solution of PBS-FC plus 0.5 mM EDTA and 0.5% bovine serum albumin (BSA) (Fisher Bioreagents, BP9704-100), and collected by centrifuging at 400xg for 3 - 5 minutes. Cell surface mycKCNQ1 was stained by incubating cells in 100 µL of PBS-FC plus 5% FBS (PBS-FC +FBS) containing anti-myc phycoerythrin (PE) (Cell Signaling, 3739S) or AlexaFluor 488-(Cell Signaling, 2279S) conjugated antibody at a 1:100 dilution for 30 minutes at room temperature in the dark. Cells were then fixed by adding 100 µL Fix & Perm Medium A (Invitrogen, GAS004) and incubating for 15 minutes. Cells were washed twice by adding 2 mL PBS-FC + FBS, centrifuging at 400 xg for 3-5 minutes, and aspirating the wash off the pelleted cells. Cells were then permeabilized and internal mycKCNQ1 was simultaneously labeled by incubating in 100 µL of Fix & Perm Medium B (Invitrogen, GAS004) with anti-myc AlexaFluor 647-conjugated antibody (Cell Signaling, 2233S) diluted 1:100 for 30 minutes. Cells were again washed 2 times with 2 mL PBS-FC + FBS and then resuspended in 400 µL PBS-FC + FBS for flow cytometry. The following controls were also prepared: untransfected (or uninduced) and unstained cells, single color controls (total protein stained with one antibody), and background staining control (uninduced or empty vector-transfected cells stained with antibodies) for use in flow cytometry analysis.

**Flow cytometry:**

Flow cytometry analysis was conducted using a 5-laser BD LSRFortessa flow cytometer (BD Biosciences, Franklin Lanes, NJ). Compensation was determined using unstained cells and single-color controls described above. Gates were drawn to select single cells that were mCherry (for LLP-int cell lines) or EGFP (for transiently transfected cells) positive. AlexaFluor647 and AlexaFluor 488 (LLP-int cell lines) or PE (transiently transfected cells) fluorescence intensities were then quantified for 2,500 cells per sample. Single cell data was exported using FlowJo data analysis software (Ashland, OR) and used for calculating average cell surface KCNQ1, total KCNQ1, and KCNQ1 trafficking efficiency. Cell surface KCNQ1 was calculated by subtracting the average background AlexaFluor 488 or PE fluorescence intensity (determined from uninduced/untransfected stained cells) from the AlexaFluor 488 or PE fluorescence intensity of each cell. Total KCNQ1 was calculated by subtracting the average background AlexaFluor 647 fluorescence intensity from the AlexaFluor 647 fluorescence intensity of each cell, multiplying this value by a brightness correction factor (to normalize for the different fluorescence intensities of each fluorophore, calculated by staining total KCNQ1 with each fluorophore and then determining the ratio of the background-subtracted fluorescence in the two samples), and then adding this internal KCNQ1 value to the cell surface value. KCNQ1 trafficking efficiency was calculated by dividing the surface value by the total value and multiplying by 100. Any background-subtracted cell surface or total values that were less than zero (below background levels of fluorescence) were recorded as “0” and any trafficking efficiency value where the surface or total levels were negative were also recorded as zero to prevent artificial calculation of high trafficking values from cells with background-level fluorescence intensities.

**RNA extraction, cDNA generation, and qPCR:**

Relative KCNQ1 mRNA levels were determined using RT-qPCR. Expression of mycKCNQ1 in the LLP-int WT mycKCNQ1 cell line was induced by treatment with 1 µg/mL doxycycline for 24 hours, then cells were treated with 0.1% DMSO or 10 µM VU0494372 for 16 hours. RNA was extracted and purified from cells using the Cold Spring Harbor “Purification of Total RNA from Mammalian Cells and Tissues” protocol (8). In short, cells were washed once with ice cold PBS, lysed directly with 1 mL per 100 mm dish surface area TRIzol (Invitrogen, 15596026), and collected by scraping with a pipette tip. The solution was collected in an Eppendorf tube, vortexed, and incubated at room temperature (RT) for 5 minutes. 0.2 mL chloroform per mL of TRIzol was added, tubes were shaken vigorously for 15 seconds, and then incubated at RT for 2 - 3 minutes. The samples were then centrifuged at 21,000 xg at 4 °C for 10 minutes. The upper aqueous phase was transferred to a fresh Eppendorf tube and 0.5 mL isopropanol per mL TRIzol was added to precipitate the RNA. After vigorous mixing and 10 minutes of incubating at RT, the tubes were centrifuged at 12,000 xg at 4 °C for 10 minutes. The supernatant was decanted off, and the RNA pellet was washed by adding 1 mL 75% ethanol per mL TRIzol, vortexing to mix, and centrifuging at 7,500xg at 4 °C for 5 minutes. The ethanol was decanted off and pellets were left at RT to dry for 5 - 10 minutes. Lastly, the dried RNA pellet was resuspended in 12 µL DEPC water, mixed by pipetting, and incubated at 42 °C for 10 minutes. RNA concentration and A260/A280 were determined with a Thermoscientific NanoDrop Lite Spectrophotometer (Waltham, MA). RNA was then reverse transcribed into cDNA using the Invitrogen SuperScript IV VILO Master Mix kit (Invitrogen, 11766050). RNA was diluted to a concentration of 125 ng/μL and 8 μL was moved to a PCR tube. 1 μL EzDNAse and 1 μL EzDNase master mix was added to each tube. Tubes were incubated at 37 °C for 2 minutes and then put on ice. 10 μL of RT master mix (2:3 RT master mix: $ddH_2O$) was added to each tube and samples were incubated at 50 °C for 10 minutes and then 85 °C for 5 minutes. Lastly, samples were diluted 1:500 in $ddH_2O$ for use in qPCR. For qPCR experiments, 2 μL diluted cDNA was combined with 18 μL qPCR master mix (10:0.4:0.4:7.2 μL 2x FAST SYBR Master mix:Fwd primer:Rev primer:$ddH_2O$) (Applied biosystems, 4385612) for each of the target genes (GAPDH – supplemental table 1, primers 17 and 18 and KCNQ1 – primers 19 and 20) in triplicate in a qPCR clear reaction plate (Applied Biosystems, A36924). Samples with no cDNA were also included for each target gene as controls. The qPCR plate was sealed and centrifuged briefly, and then qPCR was run using a CFX96 Touch Real-Time PCR Detection System (BioRad, Hercules, CA). The following program was used: 50 °C for 2 minutes, 95 °C for 10 minutes, 40 rounds of 95 °C 15 seconds and 60 °C (with detection) 1 min. Melt curves were determined by heating to 95 °C for 15 seconds, 60 °C for 1 minute and then ramping from 60 °C to 95 °C at a rate of 0.5 °C every 5 seconds with detection. $C_q$ values for each sample were calculated automatically by the instrument. Samples were analyzed by subtracting the mean $C_q$ of the KCNQ1 wells from the mean $C_q$ of the GAPDH wells for each condition ($\Delta C_q(\text{sample}) = C_q(\text{KCNQ1,sample}) - C_q(\text{GAPDH,sample})$) then subtracting the $\Delta C_q$ of the DMSO control from the $\Delta C_q$ of the drug-treated sample to get the $\Delta\Delta C_q$ for each compound. Lastly, the fold change of each sample was calculated by taking $2^{-\Delta\Delta Cq}$ for each compound treatment.

**Cycloheximide chase assay:**

WT mycKCNQ1 expression was induced in LLP-int cells by adding doxycycline to the media 24 hours before additional treatment with either 0.1% DMSO or 10 μM VU0494372 for 16 hours. To assess rate of KCNQ1 degradation, cells were then treated with either 50 μg/mL cycloheximide (CHX) (Cell Signaling, 2112S) or 0.1% DMSO (as a control) for 0, 3, 6, or 9 hours. Cells were collected and lysed, and soluble KCNQ1 was collected using the same lysis method as CETSA experiments. Total protein in lysates was quantified using Pierce 660 nm protein assay reagent (Thermo Scientific, 22660), and 6-7 μg protein was loaded per well into pre-cast NuPAGE 10% Bis-Tris SDS PAGE gels (Invitrogen, NP0303BOX) and run at a constant voltage of 100 Volts for 10 minutes, and then at 150 Volts for 60 - 75 minutes. Western blotting was performed as described for CETSA experiments, except blots were cut above the 50 kDa ladder band and the top half was probed with an anti-KCNQ1 C-terminal domain antibody (Alomone Labs, APC-022) and the bottom half was probed with an anti β-actin antibody (Cell Signaling, 8457S) as a loading control. Bands corresponding to mycKCNQ1 (approximately 70 kDa) and β-actin were quantified with the Fiji/ImageJ "band peak quantification" plugin (9, 10), and mycKCNQ1 band intensities were normalized to the β-actin intensity in the same lane. Normalized mycKCNQ1 band intensities were then represented as a "percent remaining" from the 0hr treatment time point and plotted using GraphPad Prism 10.

**Cellular thermal shift assay (CETSA):**

The effects of compounds on the thermal stability of KCNQ1 in cells was assessed with a Cellular Thermal Shift Assay (CETSA). The principles and protocol for CETSA has been described in detail in several publications (11, 12), and CETSA has also been optimized for studies of membrane proteins more broadly (13), and for studying KCNQ1 in particular (14). Briefly, LLP-int WT mycKCNQ1 cells were cultured in T-75 flasks and KCNQ1 expression was induced by treatment with 1 µg/mL doxycycline 24 hours before compound treatment. 16 hours before harvesting, cells were additionally treated with 0.2% DMSO or 15 µM VU0494372. A set of uninduced, untreated cells were also used as a control. After 16 hours of treatment, cells were washed with PBS, detached with trypsin and collected by centrifugation. Media was aspirated off the cell pellet and cells were resuspended in 1 mL PBS to wash away remaining media. Cells were centrifuged again to pellet and then resuspended in 612 µL (DMSO and VU0494372-treated samples), or 152 µL (uninduced control) PBS. A 22 µL aliquot was removed to use for counting cells. Cells were then diluted with additional PBS to bring all samples to a concentration of $1 \times 10^7$ cells/mL and protease inhibitor cocktail (Sigma-Aldrich, P8340) was added at a 1:100 dilution. Cells were then divided into 50 µL aliquots in PCR tubes. One aliquot for each condition was heated to the following temperatures: 37 °C (unheated control), 40 °C, 44 °C, 48 °C, 52 °C, 56 °C, 60 °C, 64 °C, 68 °C, 72 °C, and 76 °C for three minutes using a thermal cycler. After heat shock, 2 µL of 25% digitonin (Sigma, D141) in water was added to each PCR tube to give a final concentration of 0.96%, and each tube was subjected to three rounds of freeze-thaw lysis with liquid nitrogen. The lysate solutions were then centrifuged at 15,000 xg for 30 minutes at 4 °C. 40 µL of supernatant was transferred to a fresh tube and 1.6 µL of 0.25 M TCEP was added (final concentration 9.61 mM). Samples were incubated for 20 minutes at room temperature and then frozen at -20 °C until used for western blotting.

Soluble KCNQ1 from CETSA experiments was then quantified with western blotting. 10.5 µL of lysate was combined with 3.5 µL 4x LDS loading dye and incubated at room temperature for 20 minutes. 12 µL of each sample was loaded into a pre-cast NuPAGE 10% Bis-Tris SDS PAGE gel (Invitrogen, NP0303BOX). Samples were run at a constant voltage of 100 V for 10 minutes, and then at 150 V for 90 minutes. Proteins were transferred to a nitrocellulose membrane using the Transblot turbo system (BioRad, Hercules, CA) on the "high molecular weight" setting. After transfer, membranes were rinsed twice with dd$H_2O$ and then blocked for approximately 2 hours with blocking buffer composed of 5% BSA and 0.2% Tween-20 in tris buffered saline (TBS). Membranes were then incubated overnight shaking at 4 °C in a primary antibody solution of 1:2500 rabbit IgG anti-KCNQ1 C-terminal domain antibody (Alomone Labs, APC-022) in blocking buffer. Membranes were washed three times with TBS + 0.1% Tween-20 and then incubated in a secondary antibody solution of 1:5000 anti-rabbit IgG/HRP conjugated antibody (Cell Signaling, 7074S) in blocking buffer shaking at room temperature for 45 - 60 minutes. Membranes were again washed three times with TBS + 0.1% Tween-20. Western blots were imaged by treating with Clarity Western ECL substrate (Bio-Rad, 170-5060) and detecting chemiluminescence using an Amersham Imager 600 multipurpose imager (GE Life Sciences, Chicago, IL). Exposure times were determined automatically by the instrument and were typically in the range of 5 - 10 seconds. Bands corresponding to mycKCNQ1 (approximately 70 kDa) were quantified using Fiji/imageJ using the BandPeakQuantification plugin (9, 10). Band intensities were normalized to the intensities of the 37 °C sample band within each western blot. Data for each replicate was plotted using GraphPad Prism 10 and curve fits for each replicate were determined by using the nonlinear regression, "sigmoidal, 4PL, X is concentration" function. IC50 ($T_{Agg}$) values were calculated for each replicate from this curve fit and were plotted to compare the DMSO and compound-treated curves.

**Cell viability:**

Viability of cells after treatment with 0.1% - 0.2% DMSO or 10 - 30 µM VU0494372 was assessed with trypan blue staining. LLP-int WT mycKCNQ1 cells were cultured in 6-well plates and KCNQ1 expression was induced for ~24 hours with 1 µg/mL doxycycline. DMSO or VU0494372 was added to the cell media for ~16 hours. Cells were then detached with trypsin, retaining all media to ensure that all cells were collected. Cell solutions were diluted 1:2 in trypan blue (Invitrogen, T10282) and then cell counts and viability (percent live cells) were determined with a Countess 3 FL (Invitrogen) automated cell counter. A set of cells were set aside and killed with ~50% ethanol and then assessed for viability as a control.

**Automated patch clamp recording:**

Plasmids and site-directed mutagenesis:

Full length cDNA encoding wild type (WT) human KCNQ1 (GenBank accession number AF000571) were engineered in the mammalian expression vector pIRES2-EGFP (BD Biosciences-Clontech, Mountain View, CA, USA) as previously described (1). This vector enabled expression of untagged channel subunits with fluorescent proteins as a means for tracking successful cell transfection. The complete KCNQ1 coding region, the IRES element and fluorescent protein cDNA were sequenced in their entirety by nanopore sequencing (Primordium Laboratories/Plasmidsaurus, Arcadia, CA) and analyzed using a custom multiple sequence alignment tool (Multiple Sequence Iterative Comparator [MuSIC], available at https://doi.org/10.18131/h6hc6-n0j20). Endotoxin-free plasmid DNA with correct sequence was purified (Nucleobond Xtra Maxi EF, Macherey-Nagel Inc.), and re-suspended in endotoxin-free water.

Electroporation:

Plasmid encoding wild-type (WT) or mutant KCNQ1 was transiently transfected into Chinese hamster Ovary (CHO) cells stably transfected with human KCNE1 (CHO-KCNQ1 cells) by electroporation using the Maxcyte STX system (MaxCyte Inc., Gaithersburg, MD, USA) to generate WT $I_{Ks}$-expressing cells, as previously described (1). CHO-KCNE1 cells grown to 80-90% confluence were harvested using 0.25% trypsin (Thermo Fisher Scientific, Waltham, MA). A 500 µL aliquot of cell suspension was then used to determine cell number and viability with an automated cell counter (ViCell, Beckman Coulter, Brea, CA, USA). Remaining cells were collected by gentle centrifugation (160 × g, 4 minutes), washed with 5 mL electroporation buffer (MaxCyte Inc., EBR100) and re-suspended in electroporation buffer at a density of $10^8$ viable cells/mL. To a 400 µL cell suspension 15 µg of WT KCNQ1 cDNA were added and the DNA-cell suspension was then transferred to an OC-400 processing assembly (MaxCyte Inc.) and electroporated using the “CHO-PE” preset protocol. Immediately after electroporation, 10 µL of recombinant human deoxyribonuclease I (dornase alpha, Genentech, Inc., San Francisco, CA) was added to the DNA-cell suspension and the entire mixture was transferred to a 35 mm tissue culture dish and incubated for 30 min at 37 °C in 5% CO2. Following incubation, cells were gently re-suspended in culture media, transferred to 2 T-150 tissue culture flasks, and grown for 24 hours at 37 °C in 5% CO2. Following incubation, cells were harvested, counted, transfection efficiency determined by flow cytometry (see below) and then frozen in 1 mL aliquots at $1.8\times10^6$ viable cells/mL were stored in liquid nitrogen until they were used in experiments.

Transfection efficiency was evaluated before freezing using a benchtop flow cytometer (CytoFLEX, Beckman Coulter). Forward scatter (FSC), side scatter (SSC), and green fluorescence (FITC) were recorded. FSC and SSC were used to gate single viable cells and to eliminate doublets, dead cells and debris. Ten thousand events were recorded for each sample. Non-electroporated cells were assayed as a control for all parameters and used to set the gates for each experiment. A 488 nm laser was used to excite FITC, and the percentage of fluorescent cells was determined from the gated population.

Data were analyzed and plotted using DataController384 V1.8 (Nanion Technologies), Excel (Microsoft Office 2013), SigmaPlot 2000 (Systat Software, Inc.) and Prism 8 (GraphPad Software) software packages as previously described (1). Whole-cell currents were normalized for membrane capacitance and results expressed as mean ± 95% confidence interval (95% CI). Peak currents were recorded at 1990 ms after the start of the voltage pulse, while tail currents were measured 10 ms after changing the membrane potential to -30 mV. The voltage-dependence of activation was calculated by fitting the normalized G-V curves with a Boltzmann function (tail currents measured at -30 mV). Effects of VU0494372 are presented relative to the control (DMSO) channel assayed in parallel as percent current density measured at +60 mV and difference (Δ) in voltage-dependence of activation $V_{½}$. Statistical analysis was performed using t-test (2 conditions) or one-way analysis of variance (3 or more conditions).

**Computational binding site prediction of VU0494372:**

Computational docking was used to identify potential binding sites of VU0494372 to KCNQ1. An initial conformer of VU0494372 was obtained in Structured Data Files (SDF) format from the Vanderbilt Discovery Collection compound library managed by the Vanderbilt High Throughput Screening Core. To avoid *a priori* assumptions about binding pocket or docking pose, VU0494372 was first blind docked to two different KCNQ1 conformations from published structural models: PDB 8SIK (15), which represents the voltage sensor up conformational state with pore domain closed, and PDB 7XNK (16), which represents the voltage sensor active state with pore domain open. For the input protein structures used for docking, calmodulin and any ions were removed from both PDBs and ML277 was removed from PDB 7XNK. Only the KCNQ1 tetramer was kept, including transmembrane and cytosolic domains. We separately blind docked VU0494372 to each KCNQ1 construct using DiffDock, a diffusion generative model which outperforms traditional and deep learning docking methods in blind docking benchmarks (17). DiffDock outputs ten binding poses per ligand and protein structure combination, ranked by their predicted confidence scores. The confidence score indicates the quality of the top-ranked pose and empirically correlates with binding affinity. Seven of the ten predicted poses generated by DiffDock using 8SIK place VU0494372 at a site partially overlapping with the known docking site of ML277. We verified this was not an inherent bias of DiffDock by comparing the docked poses of other compounds from the VU library. All of the predicted poses generated by DiffDock using 7XNK place VU0494372 inside the pore domain, likely due to the large pocket formed in the open state of the pore. Using the docked poses generated by DiffDock with PDB 8SIK as starting positions, we applied a thorough targeted docking procedure to refine and score the poses. We used the BioChemicalLibrary ConformerGenerator (18) function to perform 2000 conformer generation iterations and select the top 100 conformers, sampling conformational space and ligand flexibility. We generated a Rosetta parameters file and associated PDB files of the compound conformers. We used the RosettaLigand (19, 20) small molecule docking protocol to generate 200 bound structural models for each of the DiffDock starting positions. The RosettaLigand protocol allows for the ligand to translate and rotate near the starting position, then performs a high-resolution ligand and protein side chain refinement followed by protein backbone energy minimization. . We used the Score12 score function with the restore_pre_talaris_2013_behavior flag as recommended for small molecule scoring (21). RosettaLigand calculates several protein- and ligand-specific energy score terms and combines them to determine an overall docking score. Ligand-specific energy terms are calculated from a bound ligand-protein complex and an unbound model. The overall interface score is determined by the difference of the score of the bound minus unbound complex (20). A more negative ligand interface score indicates a more energetically favorable binding energy. The most energetically favorable docked poses predict interactions with residues adjacent to those in the experimentally determined ML277 binding pocket.

**Re-synthesis of VU0608712 and synthesis of analogs**

Materials:

Solvents were obtained from commercial sources. Commercial reagents were used as received.

Instrumentation:

The compounds were purified by Teledyne ISCO normal phase column chromatograph system. Where necessary preparative reverse phase HPLC was conducted on a Gilson HPLC system using a Phenomenex Luna column (100 Å, 50 x 21.20 mm, 5 µm C18) with UV/Vis detection. $^{1}$H NMR spectra were recorded on Bruker 400 MHz spectrometer and are reported relative to deuterated solvent signals. Data for $^{1}$H NMR spectra are reported as follows: chemical shift (δ ppm), multiplicity (s = singlet, d = doublet, t = triplet, q = quartet, quint = quintet, m = multiplet, br = broad, app = apparent), coupling constants (Hz), and integration. $^{13}$C NMR spectra were recorded on Bruker 100, 125, or 150 MHz spectrometers and are reported relative to deuterated solvent signals. LC/MS was conducted and recorded on an Agilent Technologies 6130 Quadrupole instrument. HRMS was conducted and recorded at the Notre Dame Mass Spectrometry and Proteomics Facility on a micrOTOF II instrument.

2-((4-(2,4,4-trimethylpentan-2-yl)phenoxy)methyl)oxirane

A solution of NaH (388 mg, 60% Wt, 9.69 mmol) and 4-(2,4,4-trimethylpentan-2-yl)phenol (1.0 g, 4.85 mmol) in anhydrous DMF (8.0 mL) was cooled to 0°C under an argon atmosphere. After 15 min at 0°C, 2-(chloromethyl)oxirane (1.12 g, 12.1 mmol) was added. This solution was allowed to slowly warm to R.T. then it was heated at 50 °C and allowed to stir for 16 h. At this time the reaction was allowed to cool to R.T. and concentrated *in vacuo.* The resulting residue was diluted with EtOAc, washed with $H_2O$ dried with $MgSO_4$ and concentrated *in vacuo.* The crude product was purified by ISCO column chromatography eluting with 0 to 20% EtOAc in Hexane to afford the product (902 mg, 71%); $^1$H NMR (400 MHz, $CDCl_3$) δ 7.30 (d, *J* = 8.7 Hz, 2H), 6.85 (d, *J* = 8.7 Hz, 2H), 4.20 (dd, *J* = 11.0, 3.4 Hz, 1H), 3.99 (dd, *J* = 11.0, 5.4 Hz, 1H), 3.39-3.35 (m, 1H), 2.92 (dd, *J* = 4.8, 4.2 Hz, 1H), 2.77 (dd, *J* = 4.9, 2.6 Hz, 1H), 1.72 (s, 2H), 1.36 (s, 6H), 0.73 (s, 9H). LCMS ESI-MS(*m/z*) calc'd for $C_{17}H_{27}O_2$ $[M+H]^+$ 263.19, Measured 263.2.

1-(4-methylpiperazin-1-yl)-3-(4-(2,4,4-trimethylpentan-2-yl)phenoxy)propan-2-ol (VU0494372)

To a solution of 2-((4-(2,4,4-trimethylpentan-2-yl)phenoxy)methyl)oxirane (50.0 mg, 0.191 mmol), in MeOH (1.5 mL) was added 1-methylpiperazine (21.0 mg, 0.210 mmol). This was placed in an Argon atmosphere and allowed to heat at 65°C for 16 h. At this time the reaction was cooled to R.T. and concentrated *in vacuo.* The crude product was purified by ISCO column chromatography eluting with 0 to 60% MeOH in DCM to afford VU0494372 (44.5 mg, 64%); $^1$H NMR (400 MHz, $CDCl_3$) δ 7.22 (d, *J* = 8.8 Hz, 2H), 6.84 (d, *J* = 8.8 Hz, 2H), 4.15-4.07 (m 1H), 3.97 (d, *J* = 4.9 Hz, 2H), 3.49 (bs, 2H), 2.73 (bs, 2H), 2.61-2.46 (m, 6H), 2.32 (s, 3H), 1.71 (s, 2H), 1.35 (s, 6H), 0.72 (s, 9H). $^{13}$C NMR (100 MHz, $CDCl_3$) δ 156.49, 142.70, 127.20, 113.86, 70.35, 65.73, 60.64, 57.14, 55.23, 46.07, 38.09, 32.46, 31.90, 31.81. HRMS ESI-MS(*m/z*) calc'd for $C_{22}H_{39}N_2O_2$ $[M+H]^+$ 363.3006, Measured 363.3006.

1-morpholino-3-(4-(2,4,4-trimethylpentan-2-yl)phenoxy)propan-2-ol (VU0489336)

Synthesized in a similar manner as VU0494372 using morpholine and 2-((4-(2,4,4-trimethylpentan-2-yl)phenoxy)methyl)oxirane. VU0489336 (39.7 mg, 59%); $^1$H NMR (400 MHz, $CDCl_3$) δ 7.29 (d, *J* = 8.8 Hz, 2H), 6.85 (d, *J* = 8.8 Hz, 2H), 4.16-4.09 (m 1H), 4.00 (d, *J* = 4.9 Hz, 2H), 3.80-3.70 (m, 4H), 2.72-2.65 (m, 2H), 2.60-2.55 (m, 2H), 2.52-2.45 (m, 2H), 1.72 (s, 2H), 1.36 (s, 6H), 0.73 (s, 9H). $^{13}$C NMR (100 MHz, $CDCl_3$) δ 156.44, 142.78, 127.22, 113.86, 70.27, 67.16, 65.66, 61.30, 57.13, 53.93,

38.10, 32.46, 31.90, 31.81. HRMS ESI-MS(m/z) calc's for LCMS ESI-MS(*m/z*) calc'd for $C_{21}H_{36}NO_3$ $[M+H]^+$ 350.2690, Measured 350.2699.

<u>1-(piperidin-1-yl)-3-(4-(2,4,4-trimethylpentan-2-yl)phenoxy)propan-2-ol (VU0963991)</u>

Synthesized in a similar manner as VU0494372 using piperidine and 2-((4-(2,4,4-trimethylpentan-2-yl)phenoxy)methyl)oxirane. VU00963991 (51.2 mg, 77%); $^1$H NMR (400 MHz, $CDCl_3$) δ 7.28 (d, *J* = 8.8 Hz, 2H), 6.86 (d, *J* = 8.8 Hz, 2H), 4.13-4.06 (m 1H), 3.98 (t, *J* = 5.2 Hz, 2H), 2.67-2.59 (m, 2H), 2.52-2.47 (m, 2H), 2.43-2.35 (m, 2H), 1.72 (s, 2H), 1.65-1.57 (m, 4H), 1.52-1.45 (m, 2H), 1.36 (s, 6H), 0.73 (s, 9H). $^{13}$C NMR (100 MHz, $CDCl_3$) δ 156.59, 142.58, 127.17, 113.86, 70.55, 65.54, 61.38, 57.16, 54.87, 38.09, 32.46, 31.90, 31.81, 26.26, 24.40. HRMS ESI-MS(*m/z*) calc'd for $C_{22}H_{38}NO_2$ $[M+H]^+$ 348.2897, Measured 348.2894.

### Supplemental Figures and Tables

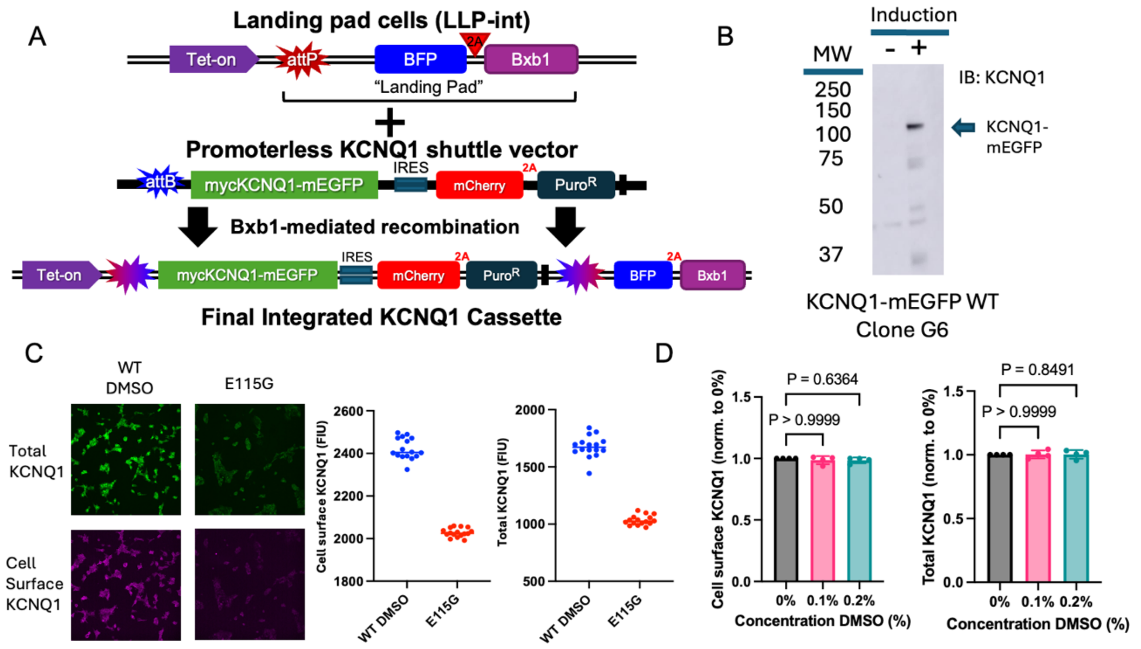


**Supplemental Figure 1. Validation of stable cell line and high-throughput screening assay**

**A)** Schematic of genomic “landing pad” site in LLP-int cells (top), target cassette in AttB shuttle vector (middle), and final integrated DNA in landing pad site (bottom). **B)** Validation of inducible expression of mycKCNQ1-mEGFP in stable cell line via western blot. Blot was probed using an anti-KCNQ1 antibody (Alomone Labs, APC-022). **C)** Representative images (left) of WT mycKCNQ1-mEGFP and E115G mycKCNQ1-mEGFP expressing cells using HTS assay labeling scheme, and representative quantifications (right) of average fluorescence intensity for DMSO-treated WT mycKCNQ1-mEGFP cells or E115G mycKCNQ1-mEGFP cells in control wells within a 384-well plate. **D)** Impact of 0.1% or 0.2% DMSO on cell surface KCNQ1 levels (left) and total KCNQ1 levels (right). Values were normalized to the 0% DMSO control in each experiment. Kruskal-Wallis test with Dunn’s multiple comparison test for follow up was used to determine P values. Trafficking efficiency values were also unchanged.

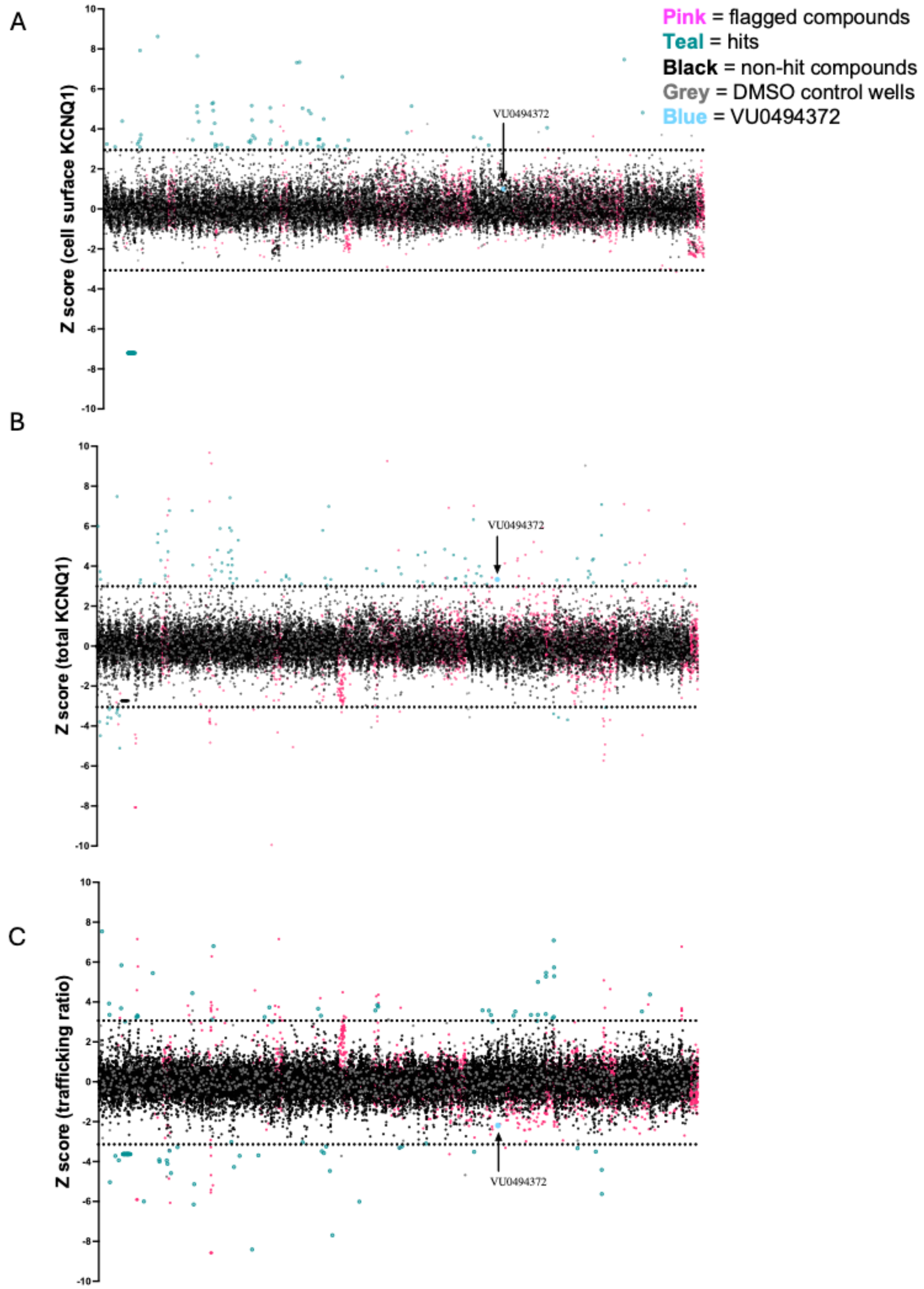


**Supplemental Figure 2. Full screen data**

Z scores for cell surface KCNQ1 **(A)**, total KCNQ1 **(B)** and trafficking ratio **(C)** for 23,611 screened compounds. Black = non-hit compounds. Teal = hit compounds. Pink = compounds flagged for poor focus and/or toxicity. Grey = DMSO control wells. Blue = VU0494372. Dotted lines indicate Z scores of +3 and -3. VU0494372 is highlighted in pink in each plot.

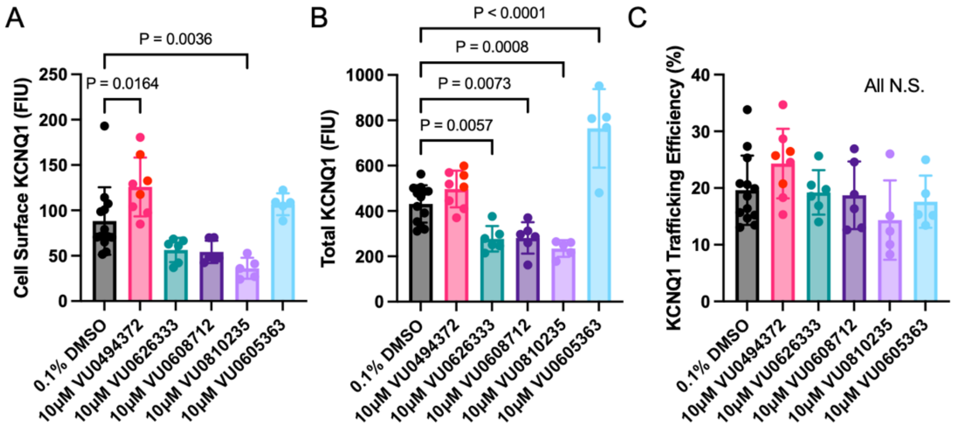


**Supplemental Figure 3: Top five hit compounds**

Quantifications of cell surface KCNQ1 **(A)**, total KCNQ1 **(B)**, and KCNQ1 trafficking efficiency **(C)** for WT mycKCNQ1 expressed in LLP-int cells after 16-hour treatment with 0.1% DMSO as a control or 10 µM compound. N=5-13. P values were determined with one-way ANOVA with Dunnett's multiple comparisons test for follow-up.

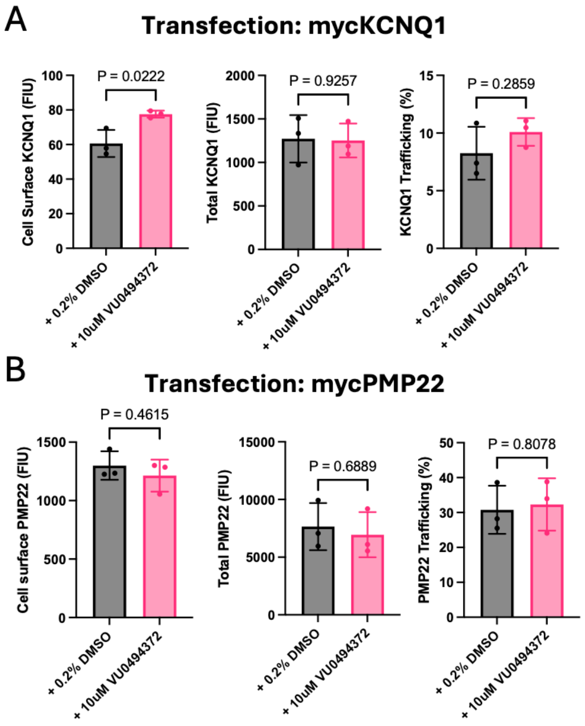


**Supplemental Figure 4: VU0494372 does not impact PMP22 expression or trafficking**

HEK 293 cells transiently transfected with KCNQ1 **(A)** or PMP22 **(B)**. 24 hours after transfection, cells were treated with 10 µM VU0494372 or 0.1% DMSO. Cells were incubated with compound or DMSO for 16 hours and surface (left) and total levels (middle), as well as trafficking efficiency (right), of KCNQ1 and PMP22 were calculated using a flow cytometry-based trafficking assay. N=3. P values were determined with unpaired t-tests.

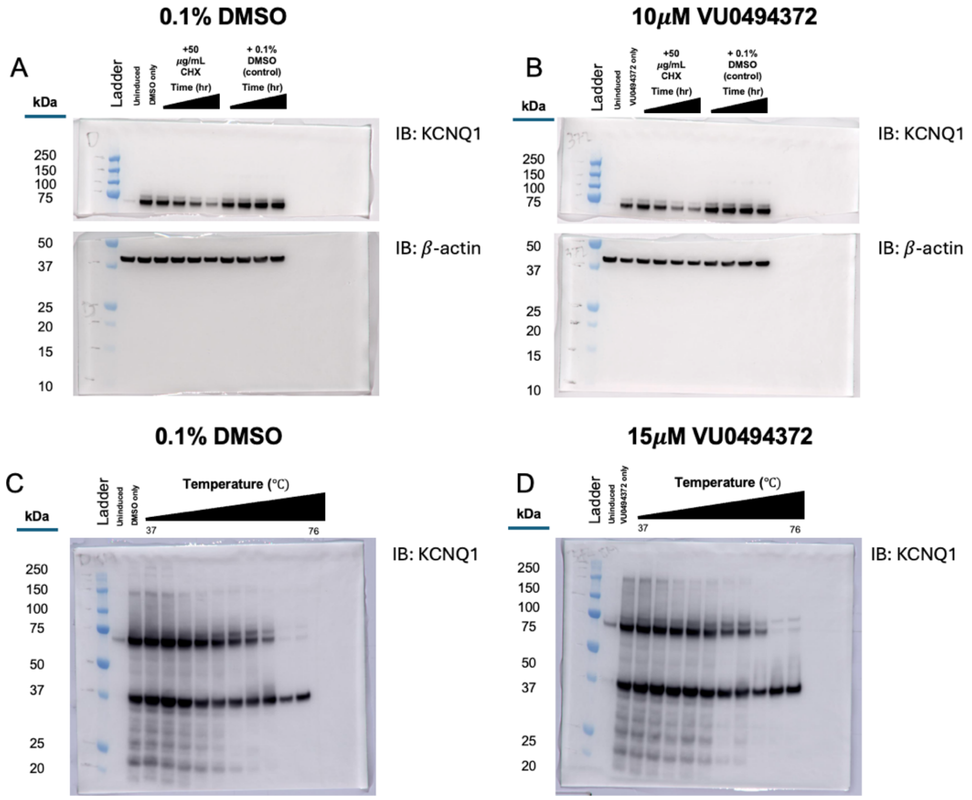


**Supplemental Figure 5: Representative blots for cycloheximide chase assays and CETSA**

Top: representative western blots from cycloheximide chase assays. Blots were cut just above the 50 kDa molecular weight marker. The top half of a blot was probed with a rabbit anti-KCNQ1 primary antibody (Alomone Labs, APC-022), and bottom half of the blot was probed with a rabbit anti-$\beta$-actin primary antibody (Cell Signaling, 8457S). Both halves were probed with an anti-rabbit HRP-conjugated secondary antibody (Cell Signaling, 7074S), and bands were detected with chemiluminescence. **A)** 0.1% DMSO-treated cycloheximide chase assay samples. **B)** 10 μM VU0494372-treated cycloheximide chase assay samples Bottom: representative western blots from cellular thermal shift assay (CETSA) experiments. Blots were probed with a rabbit anti-KCNQ1 primary antibody and an anti-rabbit HRP conjugated secondary antibody. **C)** 0.2% DMSO-treated CETSA samples. D) 15 μM VU0494372-treated CETSA samples.

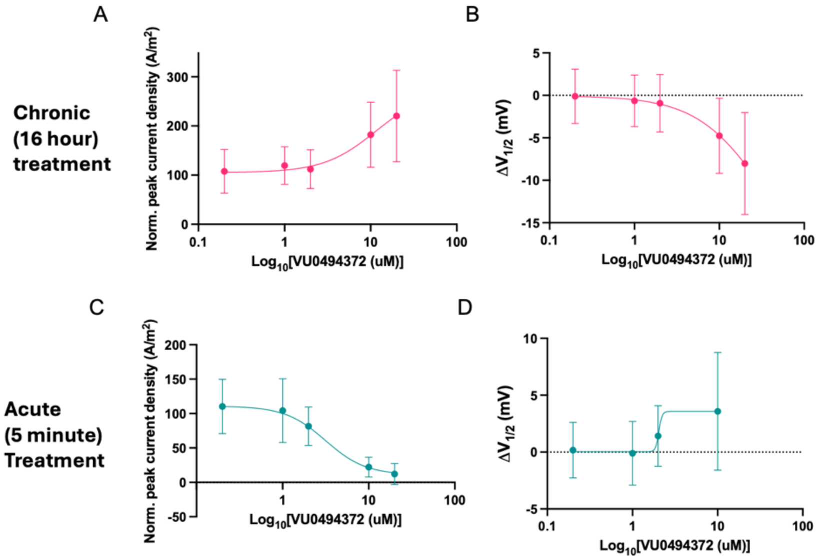


**Supplemental Figure 6: Dose-response of KCNQ1 function after VU0494372 treatment**

KCNQ1 function quantified in CHO-K1 cells stably expressing KCNE1 (CHO-KCNE1) and transfected with WT KCNQ1 using automated patch clamp electrophysiology (n=14-50 cells per group). Error bars represent standard deviation and best-fit curves were modeled in GraphPad Prism. All values were normalized to 0.2% DMSO control. Peak current density values are represented as a percent of DMSO control whereas $V_{1/2}$ values are represented as a change ($\Delta$) compared to DMSO control. A-B) Peak current density **(A)** and $\Delta V_{1/2}$ of activation **(B)** of WT KCNQ1 after 16-hour (chronic) treatment with 0.2 – 20 µM VU0494372. Compound was removed before recording (n=15-45 cells per group). C-D) Peak current density **(C)** and $\Delta V_{1/2}$ of activation **(D)** of WT KCNQ1 after 5-minute (acute) treatment with 0.2 – 20 µM VU0494372 (n=14-50 cells per group). $V_{1/2}$ of activation could not be determined at 0.2 – 20 µM VU0494372 due to low current levels.

**Supplemental Table 1.** Primer sequences used in the construction of KCNQ1 landing pad plasmids, generation of KCNQ1 variants, and qPCR experiments.

| | **Primer Name** | **Sequence** |
|---|---|---|
| 1 | Bxb_KCNQ1_F1 | GCCGGCTTGTCGACGACGGCGGTCTCCGTCGTCAGGATCATCCGCCATGGCCGCGGCCTCCTCCCCG |
| 2 | HA_Sph_KCNQ1_R | ATCTAGATCCGGTGGATCCCGCATGCTCAGGACCCCTCATCGGGGC |
| 3 | HA_SpeI_AttBxb1_F | CAACGCGGATCTTTTCTACACTAGT GCCGGCTTGTCGACGACGGCG |
| 4 | Puro_WPRE_R | AATCCAGAGGTTGATTGCGGCCGCGTGGTATGGCTGATTATGATC |
| 5 | pKanQ1_NEB_Fw | GTACAAGTAATGATCACTGTGGAATCAGAGAGACTGC |
| 6 | pKanQ1_NEB_Rev | CCTTGCTCACGGACCCCTCATCGGGGCC |
| 7 | mEGFP_NEB_Fw | TGAGGGGTCCGTGAGCAAGGGCGAGGAG |
| 8 | mEGFP_NEB_Rev | ACAGTGATCATTACTTGTACAGCTCGTCCATG |
| 9 | E115G_F | CTACAACTTCCTCGGGCGTCCCACCGGC |
| 10 | E115G_R | GCCGGTGGGACGCCCGAGGAAGTTGTAG |
| 11 | G189E_F | AGCGCAGCCGCTCCCAGAGGCC |
| 12 | G189E_R | GGCCTCTGGGAGCGGCTGCGCT |
| 13 | G179S_F | CCTCTGGTCCGCCAGCTGCCGCAGCAAG |
| 14 | G179S_R | CTTGCTGCGGCAGCTGGCGGACCAGAGG |
| 15 | V207M_F | CTCATCGTGGTCATGGCCTCCATGG |
| 16 | V207M_R | CCATGGAGGCCATGACCACGATGAG |
| 17 | GAPDH_F | TTGGCTACAGCAACAGGGTG |
| 18 | GAPDH_R | GGGGAGATTCAGTGTGGTGG |
| 19 | Q1qPCR_sense3 | GCATACTGCAGATCCTCTTCTG |
| 20 | Q1qPCr_antisense3 | TTTACCACTTCGCCGTCTTC |

**Supplemental References**

1. Vanoye CG, et al. High-Throughput Functional Evaluation of KCNQ1 Decrypts Variants of Unknown Significance. *Circ Genom Precis Med*. 2018;11(11):e002345.

2. Kanki H, et al. A Structural Requirement for Processing the Cardiac K+ Channel KCNQ1. *Journal of Biological Chemistry*. 2004;279(32):33976–33983.

3. Matreyek KA, et al. An improved platform for functional assessment of large protein libraries in mammalian cells. *Nucleic Acids Res*. 2020;48(1):e1.

4. Matreyek KA, Stephany JJ, Fowler DM. A platform for functional assessment of large variant libraries in mammalian cells. *Nucleic Acids Research*. 2017;45(11):e102–e102.

5. Shukla N, et al. Mutants of human ACE2 differentially promote SARS-CoV and SARS-CoV-2 spike mediated infection. *PLOS Pathogens*. 2021;17(7):e1009715.

6. Schlebach JP, et al. Conformational Stability and Pathogenic Misfolding of the Integral Membrane Protein PMP22. *J Am Chem Soc*. 2015;137(27):8758–8768.

7. Huang H, et al. Mechanisms of KCNQ1 channel dysfunction in long QT syndrome involving voltage sensor domain mutations. *Science Advances*. 2018;4(3):eaar2631.

8. Green MR, Sambrook J. Purification of Total RNA from Mammalian Cells and Tissues. *Cold Spring Harb Protoc*. 2020;2020(1):pdb.prot101659.

9. Schindelin J, et al. Fiji: an open-source platform for biological-image analysis. *Nat Methods*. 2012;9(7):676–682.

10. Ohgane K, Yoshioka H. Quantification of Gel Bands by an Image J Macro, Band/Peak Quantification Tool v1 [preprint]. 2019. https://doi.org/10.17504/protocols.io.7vghn3w.

11. Jafari R, et al. The cellular thermal shift assay for evaluating drug target interactions in cells. *Nat Protoc*. 2014;9(9):2100–2122.

12. Martinez Molina D, Nordlund P. The Cellular Thermal Shift Assay: A Novel Biophysical Assay for In Situ Drug Target Engagement and Mechanistic Biomarker Studies. *Annu Rev Pharmacol Toxicol*. 2016;56(1):141–161.

13. Kawatkar A, et al. CETSA beyond Soluble Targets: a Broad Application to Multipass Transmembrane Proteins. *ACS Chem Biol*. 2019;14(9):1913–1920.

14. Brewer KR, et al. Integrative analysis of KCNQ1 variants reveals molecular mechanisms of type 1 long QT syndrome pathogenesis. *Proc Natl Acad Sci USA*. 2025;122(8):e2412971122.

15. Mandala VS, MacKinnon R. The membrane electric field regulates the PIP $_2$ -binding site to gate the KCNQ1 channel. *Proc Natl Acad Sci USA*. 2023;120(21):e2301985120.

16. Ma D, et al. Structural mechanisms for the activation of human cardiac KCNQ1 channel by electro-mechanical coupling enhancers. *Proc Natl Acad Sci USA*. 2022;119(45):e2207067119.

17. Corso G, et al. DiffDock: Diffusion Steps, Twists, and Turns for Molecular Docking [preprint]. 2022. https://doi.org/10.48550/ARXIV.2210.01776.

18. Brown BP, et al. Introduction to the BioChemical Library (BCL): An Application-Based Open-Source Toolkit for Integrated Cheminformatics and Machine Learning in Computer-Aided Drug Discovery. *Front Pharmacol*. 2022;13:833099.

19. Meiler J, Baker D. ROSETTALIGAND: Protein–small molecule docking with full side-chain flexibility. *Proteins*. 2006;65(3):538–548.

20. Lemmon G, Meiler J. Rosetta Ligand Docking with Flexible XML Protocols. In: Baron R, ed. *Computational Drug Discovery and Design*. New York, NY: Springer New York; 2012:143–155.

21. Smith ST, Meiler J. Assessing multiple score functions in Rosetta for drug discovery. *PLoS ONE*. 2020;15(10):e0240450.
